# Gcm: a novel anti-inflammatory transcriptional cascade conserved from flies to humans

**DOI:** 10.1101/2022.05.29.493864

**Authors:** Alexia Pavlidaki, Radmila Panic, Sara Monticelli, Céline Riet, Yoshihiro Yuasa, Pierre B. Cattenoz, Brahim Nait-Oumesmar, Angela Giangrande

**Affiliations:** Institut de Génétique et de Biologie Moléculaire et Cellulaire, Illkirch, France; Centre National de la Recherche Scientifique, UMR7104, Illkirch, France; Institut National de la Santé et de la Recherche Médicale, U1258, Illkirch, France; Université de Strasbourg, Illkirch, France; Sorbonne Université, Institut du Cerveau - Paris Brain Institute - ICM, Inserm, CNRS, APHP, Hôpital de la Pitié-Salpêtrière, Paris, France

**Keywords:** immunity, neuro-inflammation, aging, transcription factors

## Abstract

Innate immunity is an ancestral process that can induce pro- and anti-inflammatory states. A major challenge is to characterise the transcriptional cascades that modulate the response to chronic and acute inflammatory challenges. The *Drosophila melanogaster* Gcm transcription factor represents an interesting candidate for its potential anti-inflammatory role. Here we explore its evolutionary conservation and its mode of action. We found that the murine ortholog *Gcm2* (*mGcm2*) is expressed upon aging, which is considered as a state of chronic inflammation. mGcm2 is found in a subpopulation of microglia, the innate immune cells of the central nervous system (CNS). Its expression is also induced by a lyso-phosphatidylcholine (LPC)-induced CNS demyelination (acute inflammation) and *mGcm2* conditional knock out mice show an increased inflammatory phenotype upon aging or LPC injection. In agreement with the role of this transcriptional cascade in inflammation, the human ortholog *hGCM2* is expressed in active demyelinating lesions of Multiple Sclerosis (MS) patients. Finally, *Drosophila gcm* expression is induced upon aging as well as during an acute inflammatory response and its overexpression decreases the inflammatory phenotype. Altogether, our data show that the inducible Gcm pathway is highly conserved from flies up to humans and represents a potential therapeutic anti-inflammatory target in the control of the inflammatory response.

## Introduction

The immune response is one of the oldest processes of living organisms. It goes from simple enzymatic reactions in bacteria to cellular and humoral pathways in more complex animals (1, 2). The main requirement for a successful immune response is the recognition of non-self through specific receptors that activate different immune pathways. The majority of both the receptors and their downstream pathways are highly conserved throughout evolution. The immune response is controlled by either pro- or anti-inflammatory cues. The pro-inflammatory JAK/STAT, Toll and NF-κB pathways are found in both insects and mammals (3–7). The TGF-β signalling pathway is associated with promotion of anti-inflammatory properties in vertebrates upon resolution of inflammation, with similar molecules being present in flies (8, 9). One of the most challenging issues is to discover transcription factors that coordinately block the inflammatory response.

Glial cells missing/Glial cell deficiency (Gcm/Glide, Gcm throughout the text) is expressed in the haemocytes of *Drosophila melanogaster*, functional orthologs of the vertebrate macrophages (10). *gcm* silencing in the fly macrophages does not on its own produce an overt phenotype, but it enhances the inflammatory phenotype triggered by the constitutive activation of the JAK/STAT pathway (11–13). While *gcm* is only expressed early and transiently in the haemocytes, its silencing also enhances the response to an acute inflammatory challenge performed well after *gcm* expression has ceased. Thus, Gcm in flies modulates the acute and chronic inflammatory responses, through mechanisms that are not fully understood.

Gcm is an atypical zinc finger transcription factor that is structurally conserved throughout evolution. The two mammalian orthologs are named *hGCM1* and *hGCM2* in humans, *mGcm1* and *mGcm2* in mice. *Gcm1* is required for the differentiation of trophoblasts in the developing placenta and its mutation is associated with pre-eclampsia (14). *Gcm2* is expressed and required mainly in the parathyroid glands, where it is necessary for the survival and the differentiation of the precursor cells but no role was described in the immune cells (15–17). Since the fly *gcm* gene, but not its orthologs, is also necessary in glia (18, 19), and since the fly glial cells constitute the immune cells of the nervous system, we speculated that this transcriptional pathway may have an anti-inflammatory role in microglia, the resident macrophages of the vertebrate nervous system. Microglia are not only responsible for the development and the homeostasis of the central nervous system (CNS) but are also involved in neuroinflammation (20–24). Moreover, recent studies suggest that the increased inflammation in the aged brain is attributed, in part, to the resident population of microglia (25). Interestingly, the heightened inflammatory profile of microglia in aging is associated with a ‘sensitised’ or ‘primed’ phenotype, which might be triggered by transcriptional pathways controlling the inflammatory response. This phenotype includes differences in morphology and gene expression of aged versus young microglia (26–30).

Here we show that the murine ortholog *mGcm2* starts being expressed in a subset of microglia upon aging. Loss of *mGcm2* enhances the aging phenotype in terms of microglia morphology and expression of pro-inflammatory markers, corroborating the hypothesis that this transcription factor has an anti-inflammatory role during chronic inflammation. Furthermore, *mGcm2* expression is induced in demyelinated lesions triggered by lysophosphatidylcholine (LPC) injection in the spinal cord and in *mGcm2* inducible conditional knock-out animals, the same challenge triggers a much stronger inflammatory reaction. Therefore, *mGcm2* is involved in chronic and acute responses. Most importantly, the human ortholog *hGCM2* is expressed in active demyelinating lesions of Multiple Sclerosis (MS) patients. Finally, chronic and acute challenges also induce the expression of *gcm* in flies. *De novo* expression of *gcm* counteracts the inflammatory phenotype, explaining its mode of action and highlighting its anti-inflammatory potential. Altogether, our data demonstrate that Gcm constitutes a highly conserved immune transcriptional cascade from flies up to humans and represents a novel potential therapeutic target in the control of the inflammatory response.

## Material and Methods

### Mouse lines

*Cx3cr1-Cre* mice were obtained from TAAM Orléans and bred to maintain them on the C57Bl/6J background. The *mGcm2^flox/flox^* line was created at the Institute Clinique de la Souris (ICS, Strasbourg) (**Figure 1A)**. Conditional knock-out (cKO) mice with their littermate controls derived from *Cx3cr1-Cre/+;mGcm2^flox/+^* males were crossed with *mGcm2^flox/flox^* females. The open field behavioural test was conducted at the ICS.

**Figure 1.**
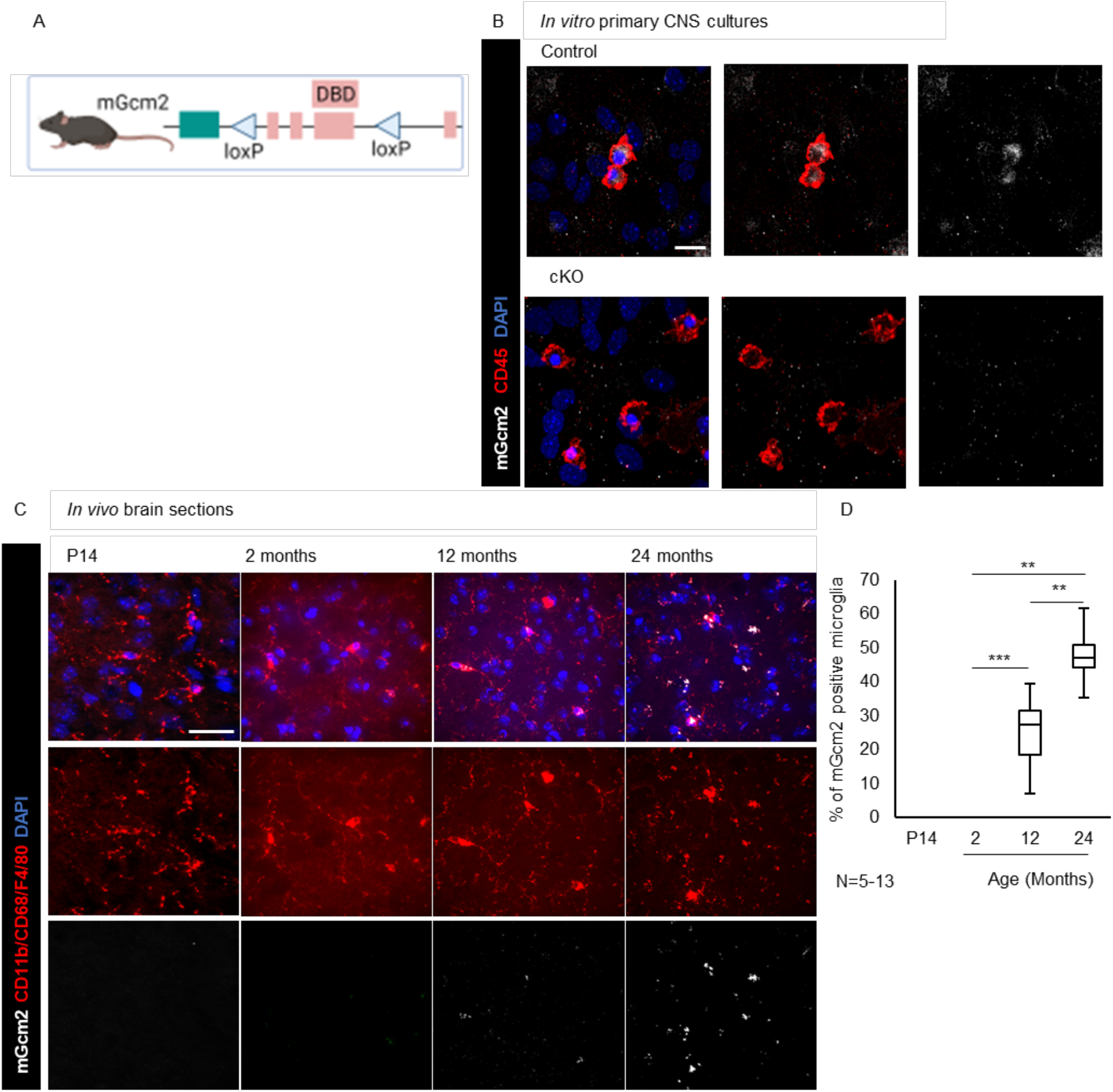
*mGcm2* locus and mGcm2 expression. (**A**) For the cKO production two loxP sites were inserted between exons 2 and 4, DBD indicates the DNA Binding Domain. (**B**) Immunolabelling of CNS primary cultures with CD45 (red), DAPI (blue) and mGcm2 (grey), N=3; scale bar 10 μm. (**C**) Immunolabelling of brain sections for mGcm2 expression *in vivo* at P14 and in 2, 12 and 24month-old animals. mGcm2 (grey), microglia (CD68, CD11b, F4/80, red) and DAPI (blue). N=5-13/genotype; scale bar: 50 μm. (**D**) Quantification of mGcm2-positive microglia in the cortex at different ages. “*” for p-value < 0.05; “**” for p-value < 0.01; “***” for p-value < 0.001. Statistical significance was determined by one-way ANOVA followed by 2-tailed, unpaired t-test. *Cx3cr1-Cre^+/-^;mGcm2^flox/+^*(control) and *Cx3cr1-Cre^+/-^, mGcm2^flox/flox^*(cKO)

To generate tamoxifen-inducible conditional knock-out *mGcm2* mice specifically in microglia, Cx3cr1^CreER/+^ knock-in mice (Parkhurst et al., 2013; JAX stock #021160) were crossed with *mGcm2^flox/flox^* mice. *Cx3cr1^CreER/+^;mGcm2^flox/+^* mice were crossed with *mGcm2^flox/flox^* animals to generate *Cx3cr1^CreER/+^*; *mGcm2^flox/flox^* mice (called icKO thereafter) and *Cx3cr1^CreER/+^; mGcm2^flox/+^* (control). WT and *mGcm2^flox/flox^* were also used as controls for demyelinating lesion experiments. All mice were maintained on a normal diet in a 12-hour light/dark cycle. 5 to 10 mice were used per group (per gender, age and genotype).

### Genotyping

DNA were extracted according to the Jacks Lab protocol. For primers, we used: *CX3Cr1^cre^* GTTCGCAAGAACCTGATGGACA and CTAGAGCCTGTTTTGCACGTTC, *mGcm2^flox^* CAATAGGGAAGTGATCCCTAGAGTC and GGGAAACTTGTCTGTTCTTTCACACAG and *mGcm2WT* CAATAGGGAAGTGATCCCTAGAGTC and GGGAAACTTGTCTGTTCTTTCACACAG, forward and reverse, respectively. Primer sequences for *Cx3cr1^CreER^* genotyping were: CX3 F-CTTCTTGCGATTCTTGCAGG; CX3 R -CACTACCTCATCATCCATGA; *CX3CreER1*- CACGGGGGAGGCAGAGGGTTT; CX3CreER2-GCGGAGCACGGGCCACATTTC.

### Tamoxifen treatment

To induce Cre recombination, adult *Cx3cr1^CreER/+^; mGcm2^flox/flox^* and *Cx3cr1^CreER/+^; mGcm2^flox/+^* mice (10 to 16week-old) were treated with tamoxifen (100mg/kg; Sigma) by intraperitoneal injections, during 5 consecutive days prior LPC-induced demyelination.

### LPC-induced demyelination of the mouse spinal cord

C57Bl6/J 12week-old females from Janvier were used for focal spinal cord demyelinated lesions. To evaluate microglial response and oligodendrocyte differentiation, we used tamoxifen treated *Cx3cr1^CreER/+^;mGcm2^flox/flox^, Cx3cr1^CreER/+^; mGcm2^flox/+^, Gcm2^flox/flox^* and WT mice. Twenty minutes before anaesthesia induction with 3% Isoflurane, animals were injected with Buprenorphine (0.1mg/kg). After induction, isoflurane concentration was increased to 2% for the surgical phase. Animals were placed on the stereotaxic frame, and small incision was made at the level of thoracic vertebrae (T8 and T9). Using a Hamilton syringe connected with a glass capillary, 1% lyso-phosphatidylcholine (LPC, sigma) was injected into the dorsal funiculus of the spinal cord. The injection site was marked with an active charcoal, and internal and external sutures were made. After surgery, animals were injected subcutaneously with Buprenorphine during 2 consecutive days and then treated *ad libitum* with a solution of Buprenorphine in the drinking water.

### P1 primary CNS cultures

Postnatal day 1 (P1) cultures were produced as previously described (31). The cultures were kept in the incubator at 37°C and 5 % CO2 for 14 days. The medium was changed at day 1 and day 3. *In vitro* cultures were fixed with 4 % PFA (Electron Microscopy Sciences) in PBS 0.1M and then proceed to the immunolabeling.

### Tissue dissections

Mice were anesthetised by a solution of Ketamine (100mg/ml)/Xylazine (Rompun, 20mg/ml): 130mg/kg of ketamine + 13mg/kg of xylazine, transcardially perfused with ice-cold PFA 4% in 0.1M PBS, and then the brains, the spinal cord, lungs and adipose tissue were dissected. The tissues were fixed overnight with 4 % PFA in 0.1M PBS and then the brains were cut into left and right hemisphere, the rest of the tissues were cut in half. Half of the samples were embedded with paraffin and the other half with cryomatrix. 8 μm and 50 μm thick sections were used for paraffin labelling and for cryo-section, respectively.

For LPC demyelinated lesions, mice were euthanised several days post LPC-injection (dpi), after lethal anaesthesia with Xylazine (10mg/kg) and pentobarbital sodium (150mg/kg), and then transcardially perfused 4% PFA in 0.1M PBS. Spinal cords were dissected and post-fixed 2 h in 4% PFA. After post-fixation, spinal cords were cryoprotected in 20% sucrose solution O/N and then frozen in O.C.T. compound (Thermo Fisher) at −60°C in isopentane. Coronal spinal cords were sections (12μm thickness) were performed at cryostat (Leica) and slides were kept on −80°C until use.

### Immunolabelling in mouse samples

For immunolabeling, samples were permeabilized with PTX (0.1M PBS, 0.1 % Triton-X100) for 30 min and incubated with blocking buffer for 1 h at room temperature (RT). The samples were incubated with primary antibodies overnight at 4°C, then incubated with the appropriate secondary antibodies. Finally, they were incubated with DAPI, (Sigma-Aldrich)) to label the nuclei and the samples were mounted with Aqua Poly/Mount (Polysciences). Primaries and their appropriate secondary antibodies are on supplementary table 1.

For peroxidase immunolabelling of Iba1, sections were incubated with the primary antibody incubation overnight and then washed extensively in 0.1M PBS,0.1% Triton. Slides were incubated for 1 h in RT with biotinylated secondary antibody for 1h, washed extensively and then incubated with the avidin–biotin–peroxidase (ABC) complex (Vector Laboratories). After washes, slides were incubated with the chromogen 3,3’ diaminobenzidine tetrahydrochloride (DAB; Sigma–Aldrich) until desired labelling intensity developed and counterstained with haematoxylin.

### Oil Red O staining

For Oil Red O (ORO) that labels macrophages containing myelin debris, spinal cord sections were dried at RT, rinsed in 60% isopropanol, then stained with freshly prepared and filtered 0.01% ORO solution. After 20 min, slides were rinsed in 60% isopropanol, counterstained with haematoxylin, rinsed in water and mount in aqueous mounting medium.

### MS tissue samples

Snap frozen post-mortem brain and cerebellar samples from MS and control patients were obtained from the UK MS tissue bank (Imperial College, London, approved by the Wales Research Ethics Committee, ref. no. 18/WA/0238). For this study, we used 4 MS and 1 control samples (**Supplementary Table 2**). 12μm-thick sections were cut on a cryostat, and lesions were classified as active (N=2), chronic active (N=1) and chronic inactive (N=2), using Luxol fast blue/major histocompatibility complex class II (MHCII) staining, as previously described (32). For immunohistochemistry, sections were post-fixed 20 min in 2% PFA and immunofluorescence labelling was performed as mentioned above for mouse experiments.

### Quantitative RT-PCR analysis

Animals were euthanised by CO2 inhalation, and samples of spinal cords around of the injection site was dissected. Total RNA, from 4 dpi LPC- and saline-injected spinal cords, were purified using RNeasy Mini Kit (Qiagen 74104). RNA concentrations were measured using Nanodrop. Reverse transcription was performed by using High-capacity cDNA reverse transcription kit with RNase inhibitor (Applied Biosystem 4374967). qPCR was performed with the TaqMan Fast Advanced Master Mix (Thermo Fisher 4444556) with specific probes for *Hprt, mGcm2, iNOS, TLR2, ARG1, Il-4ra, and CD16* (**Supplementary Table 3**). qPCR reactions with the same concentration of cDNA were run in duplicates using LightCycler 96 (Roche). *Hprt* was used as a housekeeping gene and used as endogenous control. ΔCt values were used to determine the relative gene expression change.

### RNAscope multiplex assay

RNAscope *in situ hybridisation* (RNA ISH) was performed on fixed frozen sections of mouse and human post-mortem samples. Preparations, pre-treatment and RNA ISH steps were performed according to the manufacturer’s protocols. All incubations were at 40°C and used a humidity control chamber (HybEZ oven, ACDbio). For mouse experiments, probe mixes used were as follows: *mCX3CR1* (Cat No.314221-C2), *mGcm2* (Cat No. 530481-C3), *mGcm1* (Cat No. 429661-C1), *Polr2a* (Cat No.312471, used a positive control) and *dapB (No*. EF191515; used a negative control). For RNA ISH on human tissues, probes used were: *hGCM2* (Cat.871081), *hCD68* (Cat.560591) and *Polr2a* (Cat No.310451, used as positive control); on *Drosophila*, we used the *gcm* probe (Cat No.1120751-C1)

Tyramide dye fluorophores (Cy3, Cy5: TSA plus; and Aykoya) or Opal fluorophores (Opal 520, Opal 570, Opal 690, Aykoya) were used diluted appropriately in RNAscope Multiplex TSA dilution buffer. Slides were also counterstained with DAPI.

### Imaging

Fluorescent and brightfield imaging were performed under a 20X objective using Axioscan (Zeiss) and Nanozoomer (Hammamatsu) for all quantitative analysis, and representative images in the figures were made using an Apotome (Zeiss).

Leica Spinning Disk microscope equipped with 20, 40 and 63X objectives was used to obtain confocal images with a step size of 0.2-1 μm. For the quantifications, five or more fields per sample were used with more than 50 cells in total.

### Image Analysis

Image analysis was performed with the Fiji image analysis program and with Imaris. Fiji was mainly used to produce images with sum of Z-projections. In all images, the signal was set to the same threshold in order to compare the different genotypes. Imaris (version 9.5.1) was used to analyse the morphology of microglial cells during aging using a semi-automatic protocol. The p-values were estimated after comparing control to cKO cells by two-way ANOVA test followed by bilateral student test. To analyse the activation state of microglia in LPC lesions at 7 and 14 dpi, we used the Visiopharm software. We selected 2 parameters: the number of branching points (ramifications) and roundness. Data were analysed by two-way ANOVA, followed by Tukey post hoc test.

### Transcriptome analysis of *gcmKD* haemocytes from *Drosophila* embryos and larvae

Haemocytes were sorted by FACS from stage 16 embryos of the following genotypes: *srp(hemo)Gal4/+;UAS-RFP/+* for the control and *srp(hemo)Gal4/+;UAS-RFP/UAS-gcm-RNAi* (BDRC #31519) for the *gcmKD (33*). Haemocytes were also sorted from wandering third instar larvae of the following genotypes: *srp(hemo)Gal4/HmlΔRFP* (control) and *srp(hemo)Gal4/hmlΔRFP;UAS-gcm-RNAi/+* (BDRC #31519) for the *gcmKD*. 20000 to 50000 cells were sorted for each replicate, three replicates were done for each condition. The RNA were extracted using Tri-reagent (SigmaAldrich) according to the manufacturer protocol. RNAseq libraries were prepared using the SMARTer (Takara) Low input RNA kit for Illumina sequencing. All samples were sequenced in 50-length Single-Read. At least 40×10^6^ reads were produced for each replicate. Data analysis was performed using the GalaxEast platform (http://www.galaxeast.fr/, RRID:SCR_006281) as described in (33, 34). The differential expression analysis was carried out using HTseq-Count (RRID:SCR_011867) and DESeq2 (RRID:SCR_015687) (35). The gene ontology analysis was done with ShinyGO 0.76 (36). The graphs were plotted using the packages pheatmap (RRID:SCR_016418) and ggplot2 (RRID:SCR_014601) in R (version 3.4.0) (R Core Team, 2017). The RNAseq data were deposited in the ArrayExpress database at EMBL-EBI (www.ebi.ac.uk/arrayexpress) under accession number E-MTAB-8702.

### Expression profile of the microglial markers in adult *Drosophila* haemocytes and glia

The dataset GSE79488 for adult haemocytes and GSE142788 for adult glia were retrieved from GEO database, mapped using RNA-STAR and compared with DESeq2 as described above (37). The microglial genes conserved across evolution were determined by Geirsdottir (38). The *Drosophila* orthologs were determined using DIOPT (39). The heatmap was drawn using pheatmap (RRID:SCR_016418).

### Tracing *gcm* expression

To assess the induction of *gcm* expression in *Drosophila* upon wasp infestation, a lineage tracing system was expressed under the control of *gcm*-specific promoters and a thermosensitive inhibitor. The detailed genotype is *UAS-FLP/+;act5c-FRT,y+,FRT-Gal4,UASmCD8GFP/gcmGal4,tubulinGal80^TS^;UAS-FLP,Ubi-p63E(FRT.STOP)Stinger/6KbgcmGal4 (gcm>g-trace*) for the experiment and *UAS-FLP/+;act5c-FRT,y+,FRT-Gal4,UASmCD8GFP/+;UAS-FLP,Ubi-p63E(FRT.STOP)Stinger/+* for the control (**Supplementary Figure 8A**). Embryos and larvae were raised at 18°C (tracing off) until the second instar larval stage, to avoid revealing the embryonic expression of *gcm*. Second instar larvae were infested with the parasitoid wasp *Leptopilina boulardi*. 20 female wasps were used to infest 100 *Drosophila* larvae for 2 h at RT. Infested larvae were then transferred at 29°C (tracing on) and let develop until wandering third instar larval stage for further analyses.

### Wasp infestation assays

Wasp eggs were collected upon bleeding 50 infested larvae in PBS 1X added with some N-*Phenylthiourea* crystals (Sigma) to prevent haemocyte melanisation. Wasp eggs were fixed in 4% PFA/PBS 1X for 30 min, washed with PTX (0.3% Triton X-100 in PBS 1X), incubated 1h in PTXN (NGS 5% in PTX, Vector Laboratories), incubated overnight at 4°C in primary antibodies, washed with PTX, incubated 1h in secondary antibodies, 1h with DAPI and TRITC-phalloidin (Sigma), then mounted with Vectashield mounting medium (Vector Laboratories). Haemocytes were labelled as previously published (12). Primaries and their appropriate secondary antibodies are on supplementary table 1.

For the gain and loss of function assays, we used control, gain of function (GOF) or loss of function (LOF) flies upon crossing *w;hmlΔGal4/CyoGFP;tubGal80^TS^* animals with *white-1118* (control), *UASTgcmF18A* (gain of function, GOF) or *UASgcmRNAi* (loss of function, LOF) animals, respectively (40,12). The infestation was conducted as described above and the tumour size was counted as previously described (12).

### Statistical analysis

Variance analysis using bilateral student tests for unpaired samples was used to estimate the p-values for microglia ramifications, coverage area, iNOS, Arg1-positive microglia. In each case, at least five animals were counted. In all analyses, “ns” means not significant, “*” for p-value < 0.05; “**” for p-value < 0.01; “***” for p-value < 0.001.

For cell quantification in LPC lesions, 3-5 spinal cord sections per animal were analysed, with at least N=3/group. Quantification of Olig2+ and CC1+ cells was performed using ZEN (Zeiss) and QuPath software. Statistical analyses were performed by using R and GraphPad (Prism) Software using Two-way ANOVA with Tukey post hoc comparison tests.

### Study approval

All animal experiments were conducted according to the European law for the welfare of animals. All animal procedures were reviewed and approved by the “Comités d’Ethiques en Expérimentation Animale” of IGBMC-ICS and of the Paris Brain Institute - ICM.

## Results

### Expression of the murine *Gcm* genes *in vitro*

We assessed whether the mGcm1 and mGcm2 genes are expressed in microglia by characterising CNS primary cultures (41, 42). Microglia were labelled with the broad hematopoietic marker CD45 (43). Microglia adopt one of the three morphologies both *in vitro* and *in vivo*: ramified microglia with small cell body and long ramifications, round cells with a small cell body and no ramifications, amoeboid microglia with a big cell body and no ramifications (44–46). Round cells are considered as activated microglia, the other two as resting cells (47). mGcm2, but not mGcm1, is expressed in 30% of round shaped microglia (**Figure 1B and Supplementary figure 1A**). We ascertained the specificity of the mGcm2 immunolabeling in a conditional knock out (cKO throughout the text) *mGcm2*^flox/flox^ mouse line crossed it with the *Cx3cr1-Cre* line expressed in microglia. The *mGcm2^flox/flox^* line was produced by introducing two LoxP sites upstream and downstream of the *Gcm* exons 2,3 and 4 (**Figure 1A**). mGcm2 labelling is absent in the cKO microglia, proving by the same token the specificity of the antibody and the efficiency of our cKO model (**Figure 1B).**

In sum, mGcm2 is expressed in active microglia of CNS cultures.

### *mGcm2* is expressed in the microglia of aged animals

Based on the above results, we analysed neonatal microglia at postnatal day 14 (P14) but found no mGcm2 expression (**Figure 1C**). This could be explained by the fact that, *in vivo*, microglia display major differences from microglial cultures (48). The finding that mGcm2 expression is specific to activated microglia *in vitro*, prompted us to ask whether its expression is induced in an inflammatory condition such as aging. Brain and spinal cord were collected at different ages: P14, 2, 12 and 24 months. As cell-specific markers, we used the microglia-specific cocktail (CD11b, CD68, F4/80) that distinguishes microglia from the meningeal macrophages (49). mGcm2 is expressed in microglia at 12 and 24 months but not in 2month-old animals in different areas of the brain including the cortex (**Figure 1C**). Moreover, the number of mGcm2-positive microglia increases over time: 12 and 24month-old cortices show 25% and almost double mGcm2-positive microglia (48%), respectively (**Figure 1D**).

We also performed RNA ISH on brain sections and confirmed that *mGcm2* is expressed in aged brains (18month-old animals), while *mGcm1* is not expressed in either control or cKO brains **(Supplementary figure 1B,C**). This further validates the efficiency of our flox/flox line and shows that *mGcm1* does not compensate for the lack of *mGcm2* expression. Finally, to evaluate the expression in other resident macrophages, lung and adipose tissues were dissected from 24month-old animals and cryo-sections were labelled for CD45 and mGcm2. No mGcm2 labelling was observed in the resident macrophages associated with those tissues (**Supplementary figure 1D**).

Thus, mGcm2 is expressed only in a subset of aged microglial cells.

### Microglia of *mGcm2* cKO mice have an activated morphology in homeostatic conditions

To explore the role of mGcm2 in microglia *in vivo*, we characterised their morphology, which is tightly linked to the activation state of microglia: resting cells have long ramifications while activated cells have shorter ramifications (50). Sections of control and cKO brains from different age groups were labelled with the microglial marker Iba1 (**Figure 2A-C**) (51, 52). Our criteria include the number of ramifications and the coverage area of the cell, that is, the area where microglia extend their ramifications at, which thus represents the area they can survey. The former parameter is a direct measurement of their activation state while the latter is an indirect indicator, as activated microglia have shorter or less ramification, which results in a decreased coverage area.

**Figure 2.**
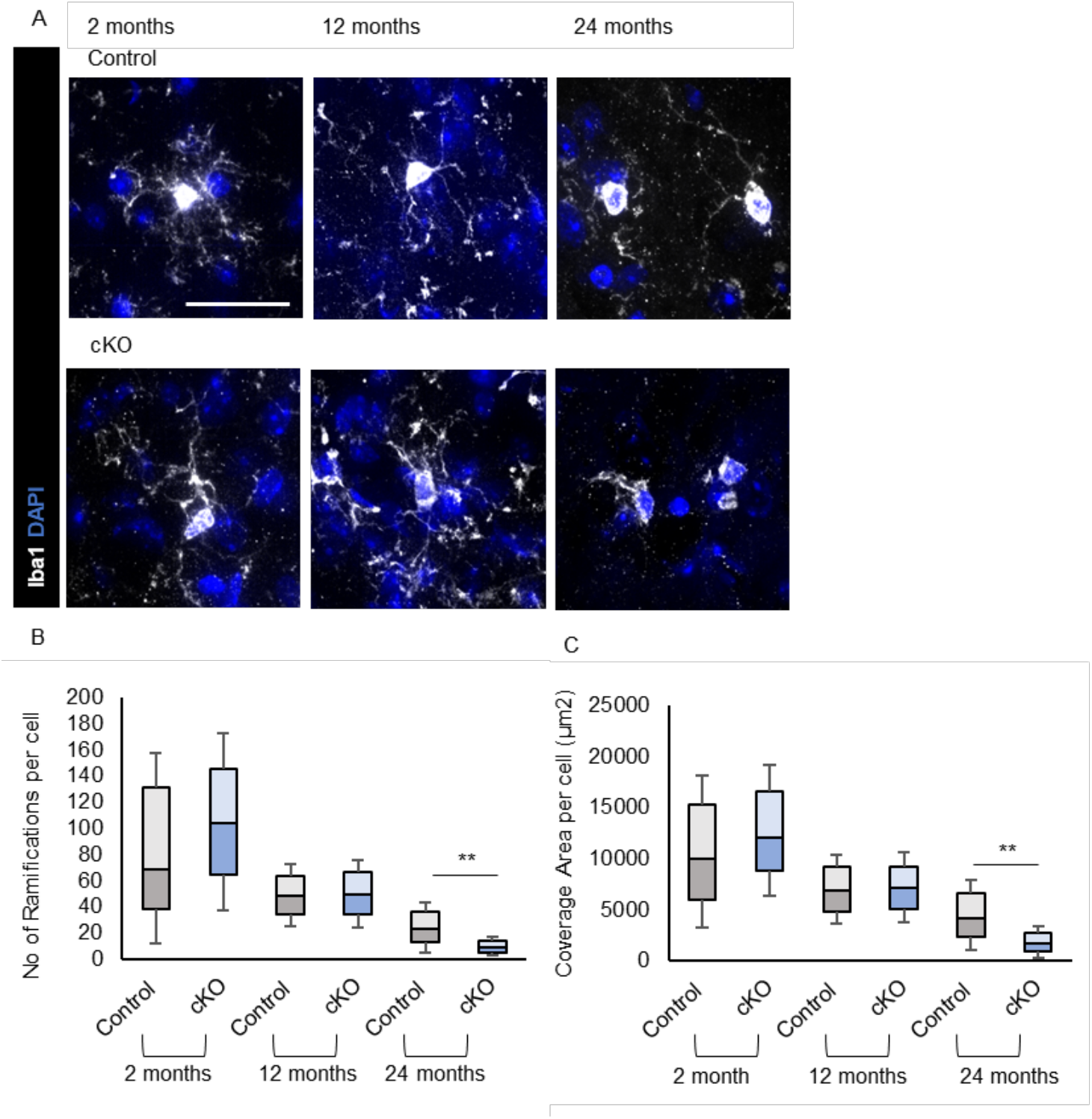
Impact of *mGcm2* deletion on microglia morphology *in vivo*. (**A**) Microglia immunolabelling of *in vivo* brain sections with Iba1 (grey) and DAPI (blue) at 2, 12 and 24 months. (**B, C**) Analysis of microglia morphology in 2, 12 and 24month-old animals of the different genotypes for number of ramifications (**B**) and coverage area (μm^2^) (**C**). N=5-13 animals per genotype; scale bar: 50 μm; p-value: *<0.05, **<0.01 and ***<0.001. Statistical significance was determined by 2-way ANOVA followed 2-tailed, unpaired t-test. *Cx3cr1-Cre^+/-^;mGcm2^flox/^* (control) and *Cx3cr1-Cre^+/-^, mGcm2^flox/flox^*(cKO)

We found that microglia morphology changes over time, but more in the cKO animals (**Figure 2A-C**). More specifically, the number of ramifications per cell decreases during aging in both genotypes (**Figure 1A**). However, there is a significant decrease in the number of ramifications in *mGcm2* cKO compared to control microglia at 24 months (**Figure 2B**). The same trend is also visible for the coverage area (**Figure 2C** tendency to decreased coverage area in the *mGcm2* cKO).

Altogether, these results reveal for the first time that the lack of *mGcm2* has an impact on microglia morphology, indicative of a pro-inflammatory phenotype.

### *mGcm2* cKO animals display a pro-inflammatory profile

We complemented the morphological data by labelling 2, 12 and 24month-old brains with pro- and anti-inflammatory markers. Microglia/macrophage activation states are classified as pro-inflammatory (or M1) or anti-inflammatory (or M2). We chose the markers inducible nitric oxide synthase (iNOS) for the M1 and Arginase-1 (Arg1) for the M2 state, which were already used to characterise microglia *in vivo* (53). Labelling from different age groups shows that *iNOS* is present mostly in 12 and 24month-old animals and its expression is increased in both groups as they age (**Figure 3A,B**). Upon quantifying the iNOS-positive microglia, we found that *mGcm2* cKO animals specifically display an increased number of iNOS-positive microglia compared to control animals by 24 months (**Figure 2B**).

**Figure 3.**
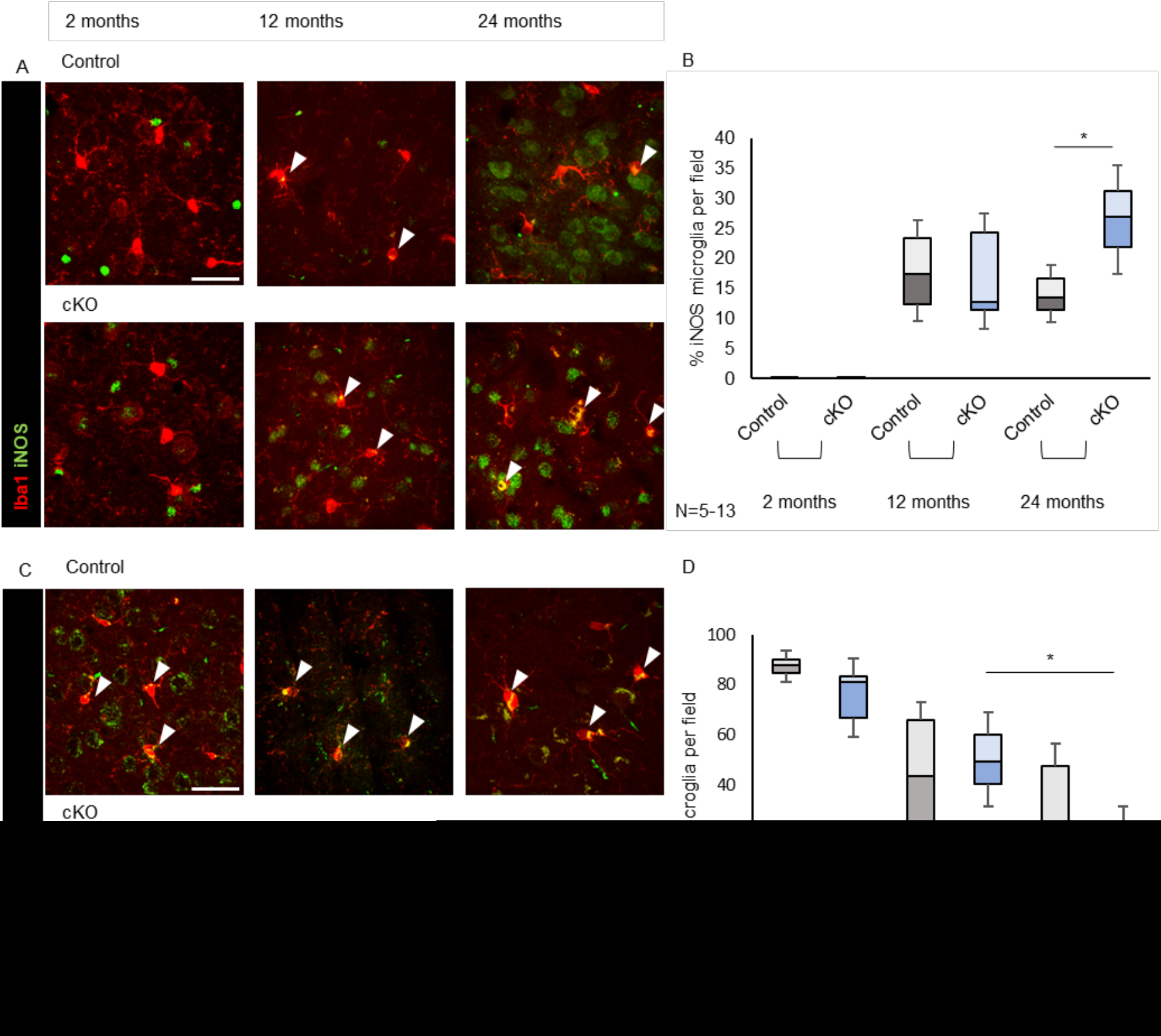
Impact of *mGcm2* deletion on microglia expression of inflammatory markers *in vivo*. (**A**) Immunolabelling of *in vivo* brain sections and (**B**) quantification of the pro-inflammatory marker iNOS (green) with Iba1 (red), (**C**) Immunolabelling of *in vivo* brain sections and (**D**) quantification of the anti-inflammatory marker Arg1 (green) with Iba1 (red), Double positive cells with Iba1 and iNOS or Arg1 expression are indicated by arrows. N=5-13; scale bar: 50 μm. p-value: *<0.05, **<0.01 and ***<0.001. Statistical significance was determined by 2-way ANOVA followed 2-tailed, unpaired t-test. *Cx3cr1-Cre^+/-^;mGcm2^flox/+^* (control) and *Cx3cr1-Cre^+/-^, mGcm2^flox/flox^*(cKO).

Next we evaluated the number of Arg1-positve cells (**Figure 3C,D**). 2month-old control animals have a higher number of Arg1-positive microglia compared to 12 and 24month-old animals (**Figure 3D**). Likewise, *mGcm2* cKO animals show a significant decrease in the number of Arg1-positive microglia, from 2 (74%) to 12 months (47%). Importantly, only the percentage of Arg1-positive microglia in the cKO animals further decreases from 12 to 24 months, resulting in a significant decline. Thus, the microglia lacking *mGcm2* display a stronger progression toward a pro-inflammatory phenotype compared to control microglia.

The state of other CNS populations does not seem overtly affected in the mutant animals. The size of astrocytes increases in inflammatory conditions (54, 55), a phenomenon called astrogliosis that can be measured via GFAP labelling (56). We found no difference between the GFAP labelling of 24month-old control and mutant animals (**Supplementary figure 2A,B**). Similarly, we found no difference in neuronal cell death by co-labelling the cell death marker caspase 3 with the pan neuronal marker NeuN (**Supplementary figure 2C-E**). No difference in oligodendrocyte number was found either (**Supplementary figure 2F,G**). Accordingly, mutant and control animals show similar behavioural habits in an open field test (**Supplementary figure 2H-J**), which assays general locomotor activity levels, anxiety, and willingness to explore.

The morphological and the molecular data show that *mGcm2* has an anti-inflammatory role in murine microglia during chronic inflammatory conditions such as aging.

### *mGcm2* is expressed in acute LPC lesions and *hGcm2* in active MS lesions

Since *mGcm2* expression is induced upon aging, we asked whether it is also expressed in immune cells following CNS injury, which represents a condition of acute inflammation. We induced acute demyelination by LPC injection in the mouse dorsal white matter spinal cord and analysed *mGcm2* expression profile. LPC lesions show limited amount of inflammation, usually associated with myelin debris removal. The mGcm2 protein was specifically detected in a subset of CD45+ immune cells (**Figure 4A**). RNAscope ISH assays also showed *mGcm2* labelling in very few microglia specifically within the lesions from 2 to 21 dpi (**Figure 4B**).

**Figure 4.**
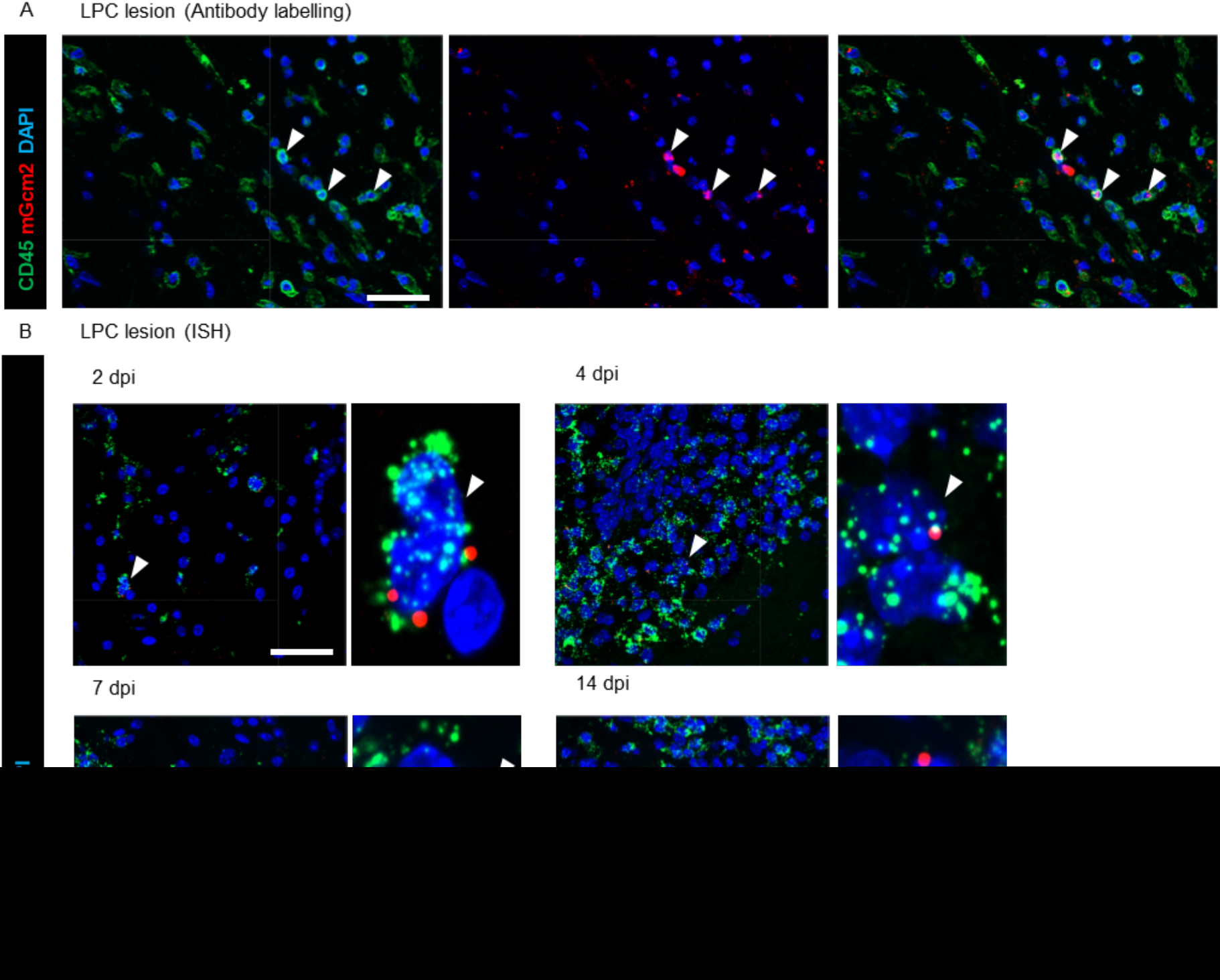
mGcm2 expression in LPC lesions of mouse spinal cord. (**A**) Immunolabelling of brain sections for CD45 (green) and mGcm2 (red) in the spinal cord at 2 dpi. Double positive cells with nuclear mGcm2 expression are indicated by arrows. (**B**) RNA ISH of brain sections, *mGcm2* (red) are mainly detected in a subset of microglial cells expressing *Cx3cr1* (green) at 2, 4, 7 and 14 dpi. Nuclei are counterstained with DAPI (blue). Scale bars: (**A**), 20μm; (**B**), 50mm.

To assess the relevance of these findings in humans, we analysed the expression of *hGCM2* in acute and chronic MS lesions, as well as in the normal appearing white matter from MS and non-neurological control cases (**Supplementary Table 1**). MS lesion subtypes were first characterised using Luxol Fast Blue and MHCII staining (**Figure 5A**) and classified as active, chronic active and chronic inactive (32). The hGCM2 protein was specifically detected in few MHCII+ immune cells located in active MS lesions, which have traditionally been defined as showing demyelination with inflammatory infiltrates, whereas chronic lesions show demyelination with little or no activity (**Figure 5B**). We next performed double RNA ISH for *hGCM2* and *hCD68* and found *hGCM2* expression in few *hCD68*-positive microglia only in active lesions and in the active rim of chronic active lesions (**Figure 5C**).

**Figure 5.**
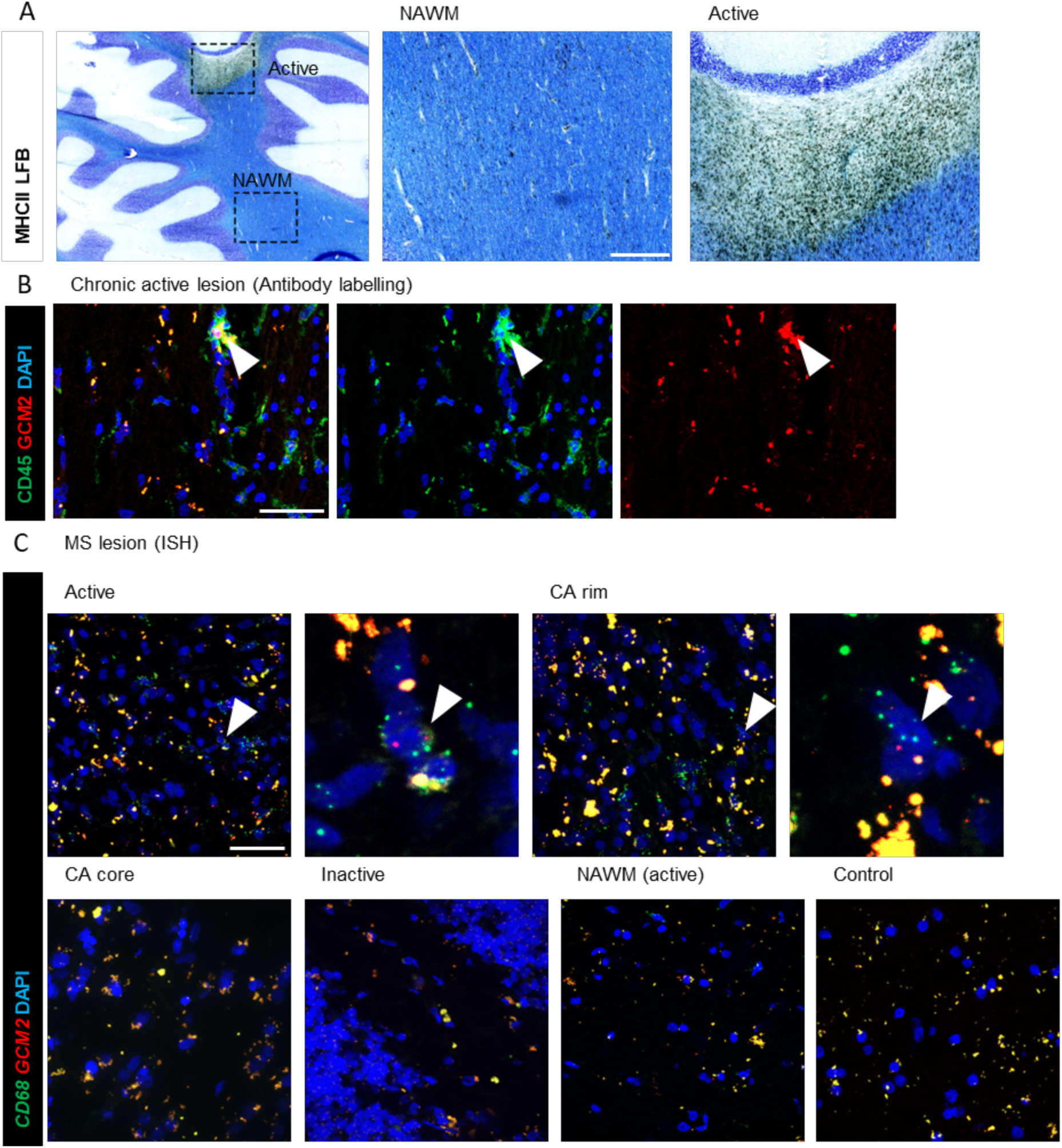
hGCM2 is expressed in microglia in active lesions of MS patients. **(A)** Luxol fast blue (LFB)/ MHCII staining showing a typical active MS lesion in the cerebellar white matter. The boxed areas illustrate an active lesion containing MHCII+ immune cells and the normal appearing white matter (NAWM) (**B**) Immunolabelling for MHCII and hGCM2 in an active lesion. Note that hGCM2 expression is only detected in few MHCII-positive immune cells (arrow) of active MS lesions. (**C**) RNA ISH for *hGCM2* and *hCD68* microglial marker in active plaques, in the active rim, chronic core of chronic active lesions, chronic inactive lesions, normal appearing white matter of MS and control cases. *hGCM2* transcripts are only detected in few *hCD68*-expressing microglial cells located in active lesions and not in inactive lesions, nor in normal appearing white matter of MS or control cases. Note that the yellow dots in panels B and C are due to lipofuscin aggregates that are inherent to human brain tissue. Scale bars: (**A**), 2mm and 500μm for magnification; (**B and C**), 50μm.

In sum, *hGCM2* and *mGcm2* gene expression is upregulated in demyelinating lesions, in a subset of cells belonging to the microglia/macrophage lineages.

### Loss of *mGcm2* in microglia promotes a pro-inflammatory response after demyelination

To decipher the functional role of *mGcm2* in microglia/macrophages in LPC lesions, we generated an inducible cKO mouse line. The *mGcm2* floxed strain (**Figure 1A**) was crossed with the *Cx3cr1^CreER/+^* mouse line (31). The deletion was induced in the F1 generation, specifically in microglia by tamoxifen injection (**Figure 6A**). We refer to this mouse strain as *Cx3cr1^CreER/+^;Gcm2^flox/flox^* (inducible cKO, icKO). *Cx3cr1^CreER/+^;Gcm2^flox/+^* heterozygous littermates (named control thereafter), *Gcm2^flox/flox^* and WT animals were used as controls for LPC lesions. 10-16 week-old mice were treated with tamoxifen during 5 consecutive days prior to LPC induced lesions. We then assessed the impact of *mGcm2* deletion in microglia on the different steps of the remyelination process, including OPC recruitment, differentiation and remyelination (**Figure 6B**). *mGcm2* deletion was first confirmed by qPCR analysis of RNA extracted from dissected spinal cord lesions at 4 dpi. *mGcm2* relative mRNA expression is indeed severely reduced in LPC demyelinated spinal cords of icKO mice with respect to controls (**Figure 6C**).

**Figure 6.**
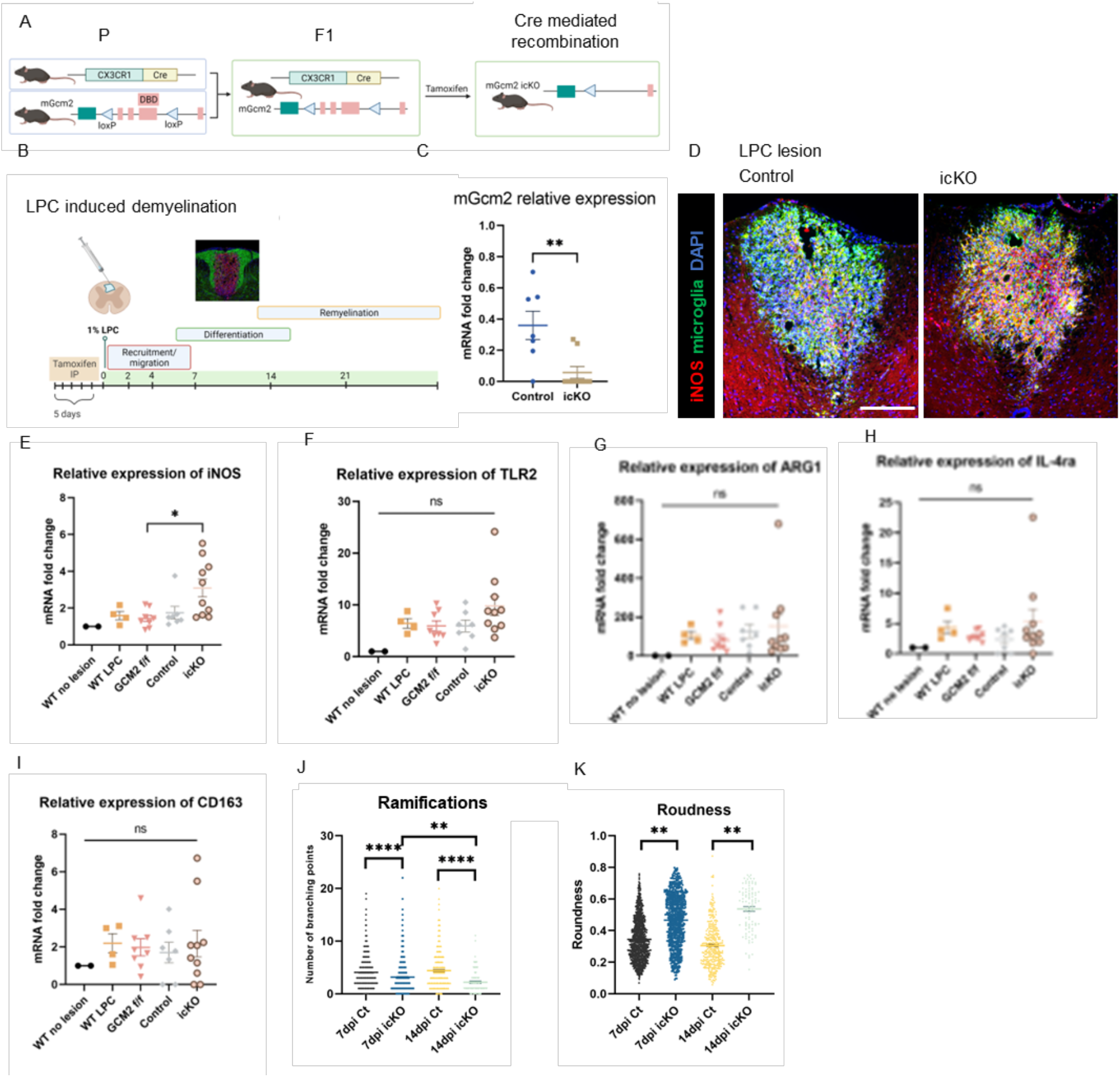
Loss of *mGcm2* function in microglial cells favours a pro-inflammatory state after demyelination. (**A**) Schematic representation of the *Cx3Cr1-Cre^ER^* transgene and *mGcm2* floxed allele and strategy used to generate tamoxifen-inducible *mGcm2* (icKO) in microglia. (**B**) Schematic illustration of LPC-induced demyelination in the mouse spinal cord dorsal funiculus and typical time frame of OPC migration/recruitment, differentiation and remyelination (**C**) *mGcm2* gene expression in demyelinated spinal cords of *Cx3cr1-Cre^ER+/-^;mGcm2^flox/+^* (icKO) and *Cx3cr1-Cre^ER+/-^;mGcm2^flox/flox^*(control) mice at 4 dpi. (C) *mGcm2* mRNA fold changes were normalised relatively to demyelinated WT spinal cords at the same time point. Data represent mean + SEM. Mann-Whitney U nonparametric test was used for the statistical analysis (N=7-8/group). **p< 0.01. (**D**) Immunolabelling for M1 marker iNOS (red) and CD11b/ F4/80/ CD68 microglia (green) in the LPC lesion of control and *mGcm2* icKO, at 14 dpi. The number of microglial cells expressing iNOS in LPC lesions increases in *mGcm2* icKO animals with respect to control. (**E-I**) qPCR analysis of the relative gene expression of M1 and M2 microglia markers in WT without LPC lesion, and LPC demyelinated spinal cords of WT, Gcm2^flox/flox^, control and *mGcm2* icKO mice. (**J-K**) Morphological analysis of Iba1-positive microglia with Visiopharm, in LPC lesions of control and *mGcm2* icKO animals, at 7 and 14 dpi. Two microglia morphological parameters were evaluated: the number of ramifications (**J**) and the roundness (**K**). Statistical analyses were performed with R and GraphPad Software using One-way ANOVA in (**E-I**) and two-way ANOVA (**J-K**), followed with Tukey post-hoc test. *p< 0.05, **p< 0.01, ****p< 0.0001. Scale bar: (**D**), 200 μm.

Once the tamoxifen-inducible Cre deletion of *mGcm2* in Cx3cr1-expressing cells had been validated, we monitored the microglia response to LPC lesions in *mGcm2* icKO mice. Immunohistochemistry for iNOS combined with the microglia cocktail markers (CD11b, F4/80 and CD68) revealed a drastic increase of microglial cells in a pro-inflammatory state in demyelinated lesions of *mGcm2* icKO mice as compared to controls (**Figure 6D**). To further confirm that the *mGcm2* deletion promotes a microglial pro-inflammatory state after LPC-induced demyelination, we performed qPCR analysis of M1 and M2 marker relative expression. In agreement with the immunohistochemistry data as well as the aging data, the expression of the M1 gene *iNOS* (**Figure 6E**) and *TLR2* (**Figure 6F**) revealed a trend increase after spinal cord demyelination in the icKO mouse strain with respect to WT, *mGcm2^flox/flox^* and control mice. We also evaluated the relative gene expression of the M2 markers *Arg1* (**Figure 6G**), *Il-4ra* (**Figure 6H**) *and CD163* (**Figure 6I**) in *mGcm2* icKO and control animals, however, our data did not reveal any significant difference.

To further corroborate these findings, we examined microglial cell morphology in conditional *mGcm2* icKO and control mice in LPC demyelinated lesions, *in vivo*. Spinal cord sections throughout the demyelinated lesions were labelled with the microglial marker Iba1 and microglia morphology was evaluated with VisioPharm (57). In line with the above data, the number of ramifications point per microglial cell decreases significantly in *mGcm2* icKO compared to controls, specifically in demyelinated lesions at 7 and 14 dpi (**Figure 6J**). As an additional morphological parameter of activated microglia, we evaluated the roundness of the cells at the same time points. The number of microglial cells exhibiting a round morphology increases significantly in *mGcm2* icKO compared to controls (**Figure 6K**), further indicating that *mGcm2* loss-of-function in microglia favours a pro-inflammatory state.

The above findings support a role of *mGcm2* as a new anti-inflammatory transcription factor in mouse microglia under pathological conditions.

### Loss of *mGcm2* in microglia delays oligodendrocyte differentiation in demyelinated lesions

Microglia activation state plays a critical role in the regulation of demyelination and remyelination (58). Therefore, we asked whether *mGcm2* loss-of-function in microglia hampers myelin debris clearance and oligodendrocyte differentiation in LPC induced demyelinated lesions. Oil-Red O staining was performed to evaluate the density of microglia containing myelin debris (**Supplementary Figure 3A**). The Oil-RedO-positive aeras as well as the percentage of Oil-RedO-positive aeras in LPC lesions do not differ significantly between *mGcm2* icKO and control mice at similar time point post-demyelination (**Supplementary Figure 3B,C**), suggesting that *mGcm2* deletion in microglia does not affect the ability of these cells to phagocyte myelin debris.

We next examined whether oligodendrocyte differentiation could be hampered. Spinal cord sections of demyelinated lesions from icKO and control mice were immunolabeled for Olig2, a pan-oligodendrocyte marker, together with CC1, a specific marker of differentiated oligodendrocytes (**Supplementary Figure 4A**). Quantification of the percentage of Olig2+CC1+ mature oligodendrocytes and Olig2+CC1-OPCs revealed a significant increase of differentiated oligodendrocytes in both groups at 7 and 21 dpi (**Supplementary Figure 4B, D**). Nevertheless, oligodendrocyte differentiation is delayed in LPC lesions of the *mGcm2* icKO strain with respect to controls, between 7 and 14 dpi (**Supplementary Figure 4B,C**). It is worth noting that the overall number of Olig2+ oligodendroglial cells in demyelinated lesions increases in both groups but was significantly lower in the icKO mice compared to the controls at 21dpi (**Supplementary Figure 4D**), suggesting that the pro-inflammatory state of *mGcm2* icKO microglia hampered oligodendroglia cell survival or apoptosis.

### *gcm* expression is induced upon aging in *Drosophila melanogaster*

Since the Gcm pathway is induced in chronic (aging) and acute (LPC induced) inflammatory conditions in mice, we asked whether this is an ancestral process and evaluated it in aging *Drosophila* brains. In physiological conditions, Gcm is expressed in a subpopulation of haemocytes, in which it has an anti-inflammatory role (12). Gcm expression is confined to the haemocytes derived from the first hematopoietic wave that occurs in the procephalic mesoderm of the embryo (19). These haemocytes cease to express Gcm by the end of embryogenesis and survive to the adult where they coexist with those derived from the second wave occurring in the larval lymph gland, which is Gcm independent (59).

In the adult, haemocytes are mostly associated with peripheral tissues (60), we therefore first assessed the number of brain associated haemocytes over time by labelling dissected fly brains with macrophage markers (P1/NimC4 and Hemese) at different ages, from week 1 (young animals) to week 6 (old animals), (**Figure 7A,B**). The number of brains that display associated haemocytes increases over time and by 6 weeks all brains are associated with haemocytes (**Figure 7C**). The number of brain-associated haemocytes observed in each animal varies, which likely also depends on the dissection protocol that may dissociate from the brain migratory cells as haemocytes. We next assessed the expression profile of Gcm in the adult brain upon aging, by performing *in situ* hybridization with a RNAscope probe. In young animals, Gcm is expressed only in two neuronal clusters located in the central brain (**Supplementary Figure 5A**) at lateral and dorsal positions (61). In 6week-old animals, however, Gcm labelling is also present at ectopic positions, indicating *de novo* expression (**Figure 7A,D**). As in the case of the haemocyte markers, *gcm* labelling is located at the surface of and not within the brain, indicative of cells that are associated with but do not belong to the tissue itself.

**Figure 7.**
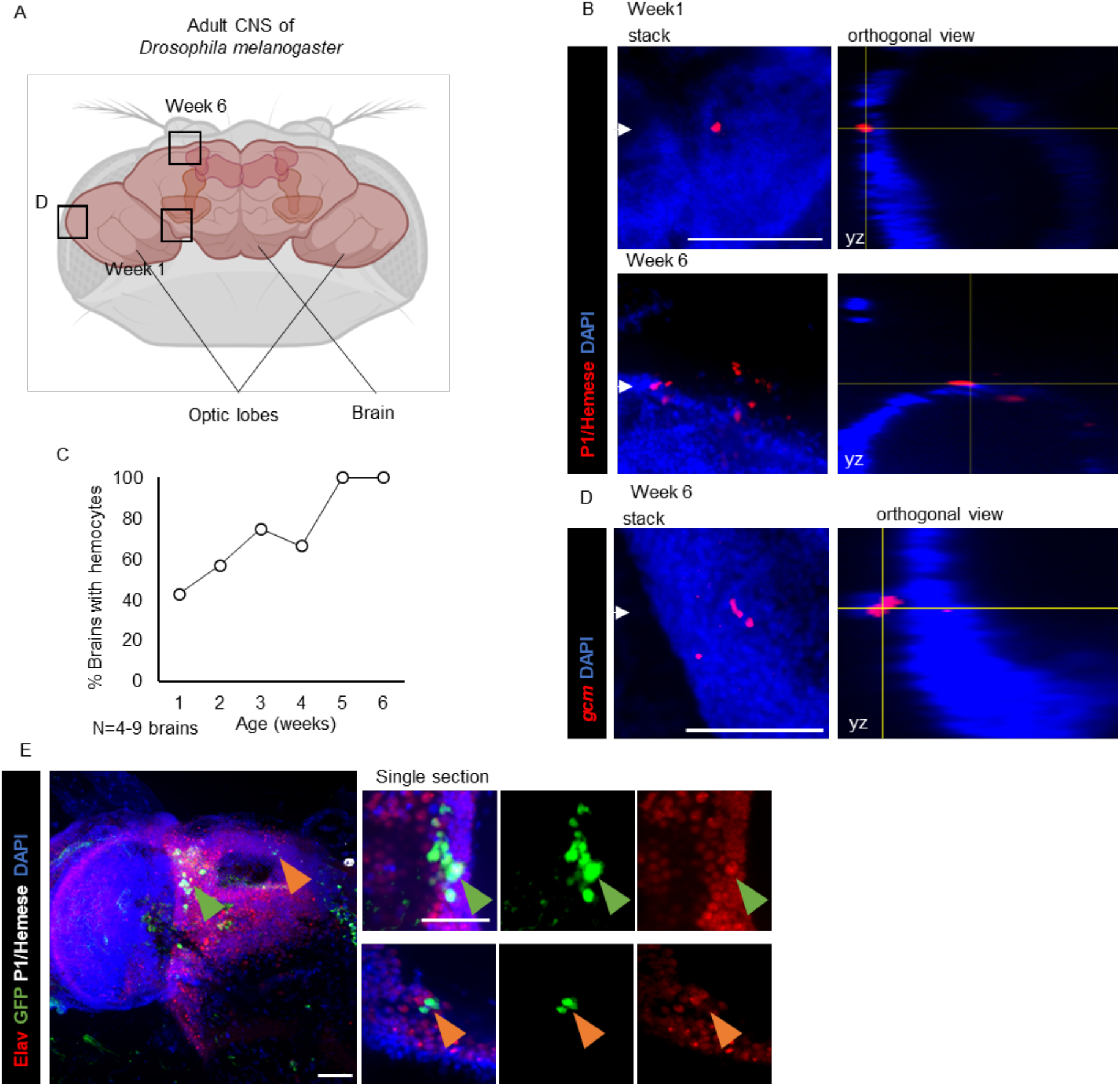
Haemocytes and *gcm* expressing cells associated with the aging *Drosophila* CNS. **(A)** Schematic of the adult *Drosophila* brain, which contains the medially located central complex and the lateral optic lobes. The squares indicate the position of the pictures (**B and D**). (**B**) Haemocytes associated with the brain at different ages, from week 1 and 6 at 25°C. The rectangular panels show an orthogonal projection along the yz axes from the position indicated by the white arrowheads. (**C**) Percentage of brains with associated haemocytes at different ages. (**D**) RNA ISH showing ectopic *gcm* expression in the aging brain, the orthogonal view shows that the signal is in cells closely associated with the brain but not within it. Scale bar: 50 μm. (**E**) Immunolabelling of adult *Drosophila* brain with *gcm* tracing (*gcm>g-trace*) in green, P1/Hemese (grey), Elav (red) and DAPI (blue), Scale bar: 20 μm

### *gcm* expression is induced upon acute challenge and counteracts the inflammatory state

*gcm* expression in the old haemocytes might be a transient process. Since the *in situ* assay only identifies cells that express *gcm* at the time of the dissection, this approach may miss cells that have expressed *gcm* earlier. For this reason, we performed lineage tracing using the g-trace tool and followed all the cells that express and/or have expressed *gcm* at some point (62). This may reveal more *gcm*-positive cells than the *in situ* assay. To make sure that we specifically look at *de novo* expression and not at remnant expression from the embryonic haemocytes, we activated the g-trace only after adult eclosion (**Figure 8A,B**). The results showed *gcm*-positive cells that are associated with the brain and do not express glial or neuronal markers. Of note, the number of *gcm*-positive cells that are associated with old brains is not higher than those revealed by *in situ*. Surprisingly, these cells do not express the P1/Hemese pan-haemocyte markers either (**Figure 7E and Supplementary Figure 5B**). This may indicate an uncharacterised population of haemocytes present at that location or these cells may have not yet acquired a proper haemocyte identity. The production of additional tools will be necessary to distinguish amongst these possibilities, but clearly non-neural *gcm* expressing cells become associated with the aged brain.

**Figure 8.**
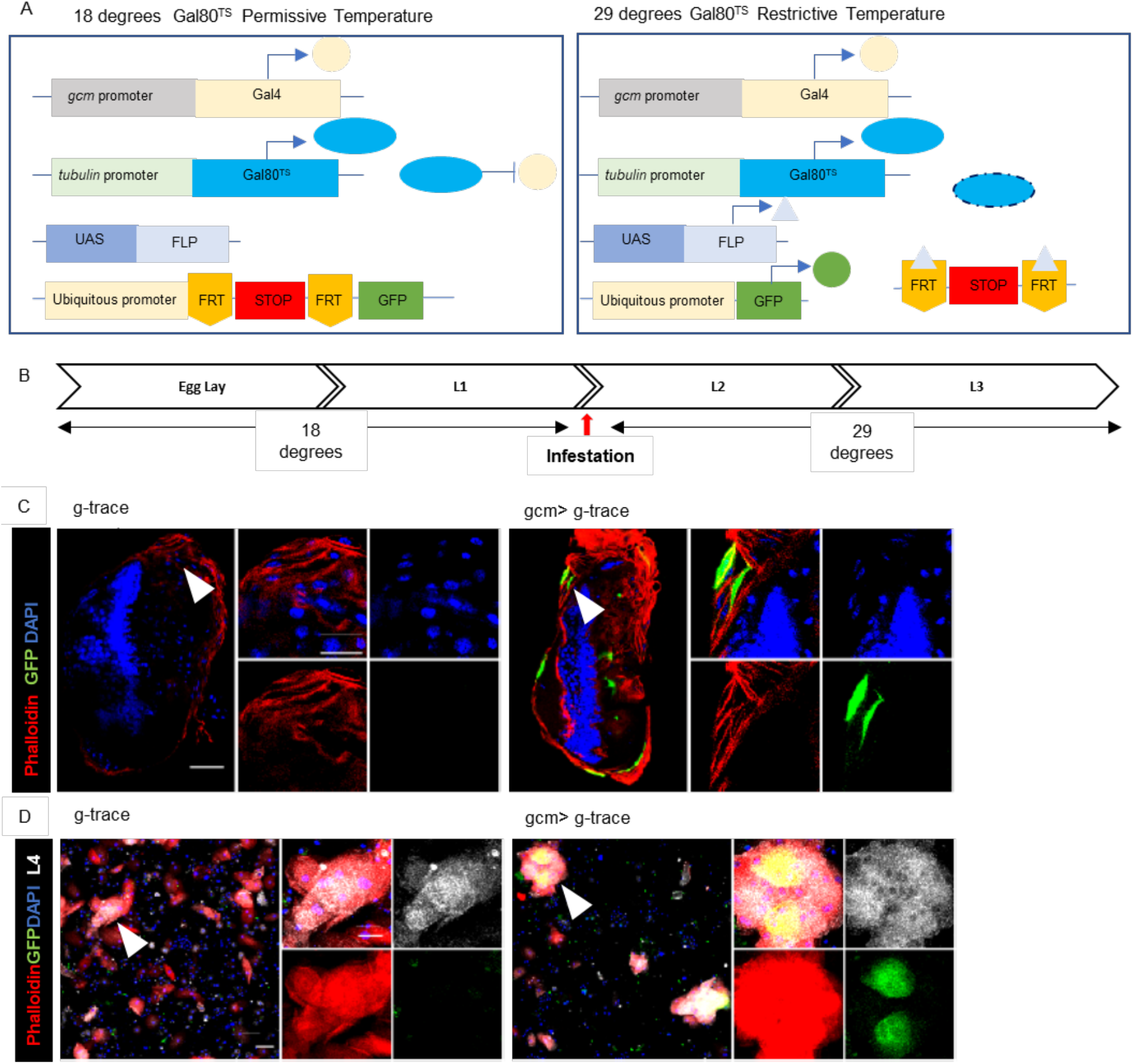
Acute inflammation induces *gcm* expression in *Drosophila*. (**A**) Inducible lineage tracing using the UAS-Gal4 system. (i) The thermosensitive Gal80 protein (Gal80^TS^) blocks Gal4 activity and hence Gal4-dependent gene transcription at 18°C. (ii) At 29°C, Gal80^TS^ is rapidly degraded and Gal4-dependent transcription can occur, FLP indicates the Flippase recombinase, FRT the Flippase Recognition Target. (**B**) Timeline of the wasp infestation assay. Larvae are kept at 18°C and upon infestation they are transferred at 29°C until the end of the larval development. (**C**) Tracing of *gcm* expression in activated haemocytes (lamellocytes) surrounding the wasp embryo (nuclei visible with DAPI (blue)) and (**D**) in haemocytes. Phalloïdin (red), *gcm>g-trace* (green), lamellocyte marker L4 (grey) and DAPI (blue). N=3; scale bar for enlarged picture: 50 μm, for magnification: 10 μm.

Based on the above results, we asked whether *gcm* is reactivated upon an acute inflammatory challenge. In *Drosophila*, the most studied acute inflammatory response is induced by wasp parasitisation. In brief, the parasitoid wasp *Leptopilina boulardi* is allowed to infest and lay eggs in *Drosophila* larvae. This leads to extensive haemocyte proliferation and activation, which consists in resting haemocytes transdifferentiating into lamellocytes (activated haemocytes) in the infested larvae (63). Lamellocytes are huge cells able to encapsulate the wasp egg, preventing it from hatching and hence allowing *Drosophila* to escape the infestation. Upon activating the g-trace tool only during the larval life, no *gcm* expression can be detected in normal conditions in third instar larvae (**Supplementary Figure 5C**). Upon wasp infestation, however, *gcm* (g-trace)-positive cells surround the wasp eggs, revealing a *de novo* expression of *gcm* following the acute challenge (**Figure 8C**). We also found *gcm* expression in circulating lamellocytes (**Figure 8D**). Thus, *gcm* expression is induced by an acute inflammatory challenge in flies.

Since *gcm* is expressed *de novo* upon wasp infestation, we asked how *gcm* gain or loss of function (GOF or LOF, respectively) would affect the response to wasp infestation (**Figure 9A**). To this purpose, we specifically induced (GOF) or silenced (LOF) *gcm* expression in the larval haemocytes <u>after</u> wasp infestation (**Figure 8B**) and evaluated the so-called tumour phenotype as a readout of the inflammatory response. The infested animals of the three genotypes (control, LOF and GOF) carry tumours, but their number and/or size varies. Large tumours contain the wasp eggs encapsulated by the fly haemocytes, while small/medium size tumours are due to haemocyte aggregations. LOF animals have in average more tumours than control and GOF animals (**Figure 9B**). This is mostly due to a very large increase in the number of small tumours (**Figure 9C,D**). GOF animals, on the other hand, show a decreased number of large tumours compared to control animals.

**Figure 9.**
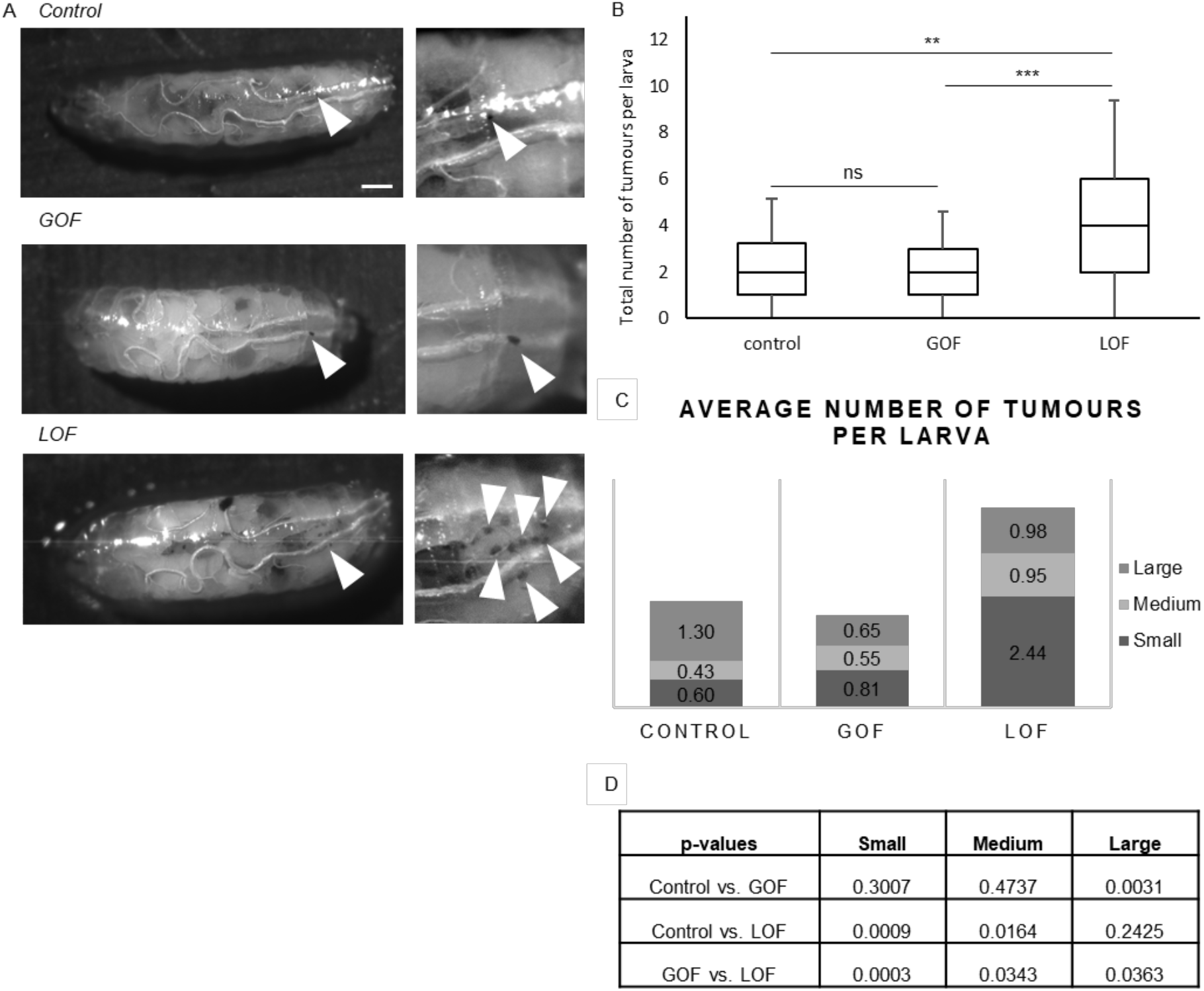
(**A**) Brightfield images of infested larvae for the control (*w;hmlΔGal4/+;tubGal80^TS^/+*), GOF (*w;hmlΔGal4/+;tubGal80^TS^/UASTgcmF18A*) and LOF (*w;hmlΔGal4/+;tubGal80^TS^/UASgcmRNAi*) genotypes.. The arrows indicate the small size tumours; scale bar 100 μm. (**B**) Quantification of the total number of tumours per larva. (**C**) Quantification of the average number of tumours per larva according to their size (**D**) and the p-values. *p< 0.05, **p< 0.01, ****p< 0.0001 and ns for not significant.

In sum, silencing the *de novo* expression of *gcm* aggravates the inflammatory phenotype and inducing *gcm* expression *de novo* ameliorates it.

### *gcm* downregulation triggers a pro-inflammatory state in the *Drosophila* haemocytes

Altogether, our data indicate that the conserved Gcm pathway is induced in response to a wide variety of challenges and counteracts acute as well as chronic inflammation. It seemingly has a priming role: the cKO mice are fully viable, fertile and do not display an overt inflammatory phenotype, much like the mutant flies (12). To investigate the molecular mechanisms underlying this priming process we proceeded to a high throughput analysis in flies, given the simplicity of this animal model. Since Gcm is also involved in gliogenesis and its mutation is embryonic lethal, we analysed the transcriptome of *srp(hemo)Gal4;UAS-gcm-RNAi (gcmKD*) animals in which Gcm is specifically affected in haemocytes. *gcmKD* haemocytes present overall increased levels of expression of immune related genes, such as anti-microbial peptides and components of major immune pathways (**Figure 10A**). In the embryo, the different expression is restricted mostly to STAT92E and Toll, but in the larva more genes appear to have different expression levels between *gcmKD* and control haemocytes. The highest difference is found in Gene Ontology (GO) terms associated with immune regulation, such as regulation of immune response to bacteria and signalling pathway of recognition of peptidoglycans (**Figure 10B)**.

**Figure 10.**
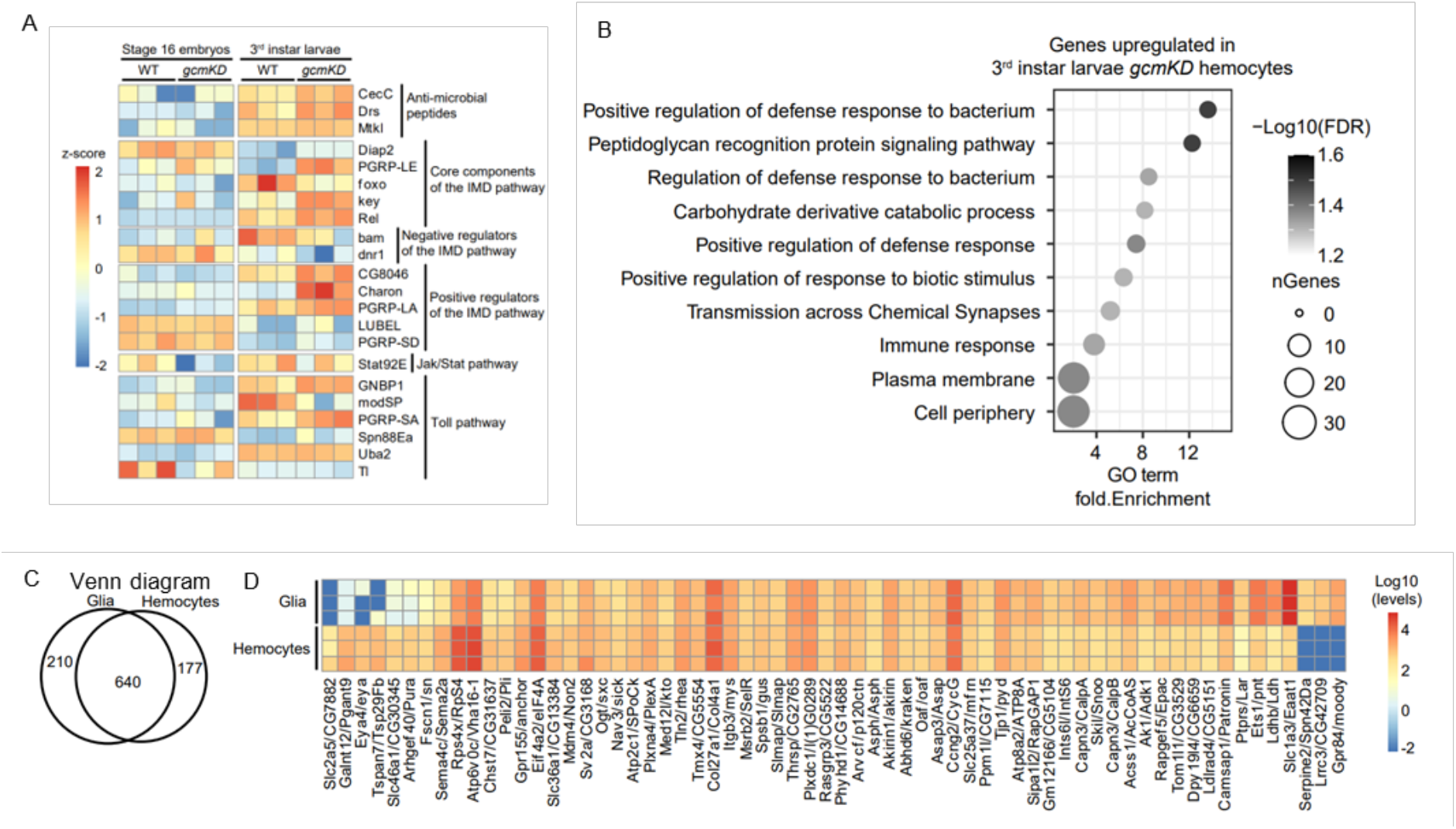
RNAseq data from microglia and from fly haemocytes and glia. (**A**) Heatmap from RNAseq data showing differentially expressed genes in heamocytes from late embryos (E16) and wandering third instar larvae. (**B**) GO term analysis of the genes upregulated in *gcmKD* haemocytes from wandering third instar larvae. (**C**) diagram representing the number of microglial markers upregulated in glia, in haemocytes or expressed in both tissues at similar levels. The genes upregulated in a specific tissue present an expression fold difference above 2 and a p-value < 0.05 (determined using DESeq2). (**D**) Heatmap representing the expression levels (Log10 scale) of the orthologs of microglial markers in *Drosophila* adult haemocytes and glia. The markers presenting the strongest expressions (>500reads) were selected. The expression levels were retrieved from the datasets GSE79488 (adult haemocytes) and GSE142788 (adult glia) and analysed using DESeq2.

Thus, silencing *gcm* triggers a pro-inflammatory state.

## Discussion

The present study identifies a novel and evolutionarily conserved anti-inflammatory transcriptional pathway. The Gcm transcription factor was known to regulate the development of fly immune cell populations (glia and haemocytes) and to modulate the inflammatory response in flies. Here we demonstrate that the expression of fly and murine *Gcm* genes is induced upon challenge, which helps counteracting the inflammatory state. Moreover, the human *Gcm2* ortholog is expressed in MS lesions. The identification of this conserved pathway opens exciting perspectives to study neuro and inflammatory diseases.

### *mGmc2* and its anti-inflammatory role in the microglia of aged mice

Aging results in gradual loss of normal function, due to changes at the cellular and molecular level (64). Microglia exhibit an exaggerated pro-inflammatory response during aging, a phenomenon referred to as microglia priming (26, 27). Morphologically, aged microglia have enlarged processes, cytoplasmic hypertrophy and a less ramified appearance (28, 65). They also express higher levels of activation markers than control microglia (66). All these changes lead to impaired remyelination and are associated with decreased number of M2 microglia, cells in an anti-inflammatory state (58).

Our data indicate that *mGcm2* expression helps keeping the inflammatory state under control during aging. Aged microglia of cKO mice have a pro-inflammatory morphology with less ramifications and coverage area. Furthermore, they show a statistically significant increase of the pro-inflammatory marker *iNOS* and a significant decrease of the anti-inflammatory marker *Arg1* between 12 and 24 months. The most parsimonious hypothesis is that *mGcm2* acts as a regulator of the activation state in microglia. Loss of its expression leads to an uncontrolled stimulation and thus a higher pro-inflammatory profile even under basal conditions. It is widely accepted that targeting anti-inflammatory (M2) regulators of microglia could lead to new therapeutic targets for aging, mGcm2 may represent one of them.

### Vertebrate *Gcm2* is a new anti-inflammatory transcription factor in CNS demyelinating lesions

The mGcm2 protein is present in few CD45-positive immune cells after acute demyelination induced by LPC injection, indicating that mGcm2 expression is restricted to a subset of inflammatory cells. Moreover, using double RNA ISH for *mGcm2* and *cx3cr1*, we provided compelling evidence supporting the expression of mGcm2 in a subset of microglia/macrophages, specifically in demyelinated lesions. Acute inflammation is one of the features of LPC-induced demyelination (67, 68). Interestingly, hGCM2 expression was also detected in a small fraction of MHCII/CD68-double positive microglia/macrophages located specifically in active lesions and in the active rim of chronic active lesions of MS, thus further supporting a conserved function of Gcm2 in microglial/macrophage lineage cells in humans. In order to evaluate the role of mGcm2 specifically during an acute inflammatory condition and not as a constitutive deletion, as we did during aging, we used the icKO model. Our data gained in the mGcm2 icKO, specifically in microglia/macrophages, clearly indicate that mGcm2 favours an anti-inflammatory M2 state, and could therefore promote indirectly myelin regeneration and repair by modulating OPC differentiation after acute demyelination (58). This emerging anti-inflammatory role of Gcm2 may open new therapeutic interventions, targeting this transcription factor in demyelinating diseases.

### Gcm: glia, haemocytes and microglia

Our data explain what was considered as a conundrum. The fly *gcm* gene was initially described for its developmental role in glia and haemocytes (10, 18). Surprisingly, while the genes necessary for neuronal differentiation are functionally conserved in evolution, the orthologs of Gcm, the fly glial promoting factor, are neither expressed nor required in the differentiation of vertebrate glial cells (69, 70). Moreover, none of the early transcription factors expressed/required in fly glial cells and target of Gcm such as the panglial Reverse Polarity (Repo) transcription factor is expressed in vertebrate glia and this gene is not even conserved in the vertebrate genome. The gliogenic pathway seems therefore not evolutionarily conserved, in line with the hypothesis that glia may have evolved several times, adapting to the needs of the organism (71). The glia of simple organisms play a neural role, as they control axon ensheathment, synapse activity and insulate the brain through the blood brain barrier (BBB). In addition, they act as the resident immune cells of the nervous system. By contrast, in complex organisms, the role of brain resident immune cells is taken by a new cell type of non-neural origin, the microglia, that infiltrate the nervous system before the BBB is formed. This division of labour guarantees a better and more targeted response to inflammatory challenges. We speculate that the haemocytes of simple, fast developing, animals migrate along the nervous system and contribute to remove dying cells, but as the BBB forms, they are excluded from the tissue, the immune function being taken up by the glial cells. According to this hypothesis, we should find orthologs of microglial markers expressed in fly haemocytes as well as glia. A recent and elegant study characterised the microglia transcriptional programs across ten species spanning more than 450 million years of evolution (72). By using these data with the RNAseq data from adult fly haemocytes and glia, we created a heatmap of the microglia orthologs in *Drosophila* (37, 73). These orthologs are mostly shared between the two fly cell populations (**Figure 10C,D**). One microglial gene whose ortholog is shared by haemocytes and glia is Peli2, a member of E3 Ubiquitin ligases controlling the Toll signalling pathway (74, 75). The *Pellino (Pli*) fly gene antagonises Toll-mediated innate immune signalling by controlling MyD88 turnover in macrophages. Future studies will determine whether the pathway is also conserved in glia. In addition to the shared genes, some are specific to glia (*spn42Da, moody, CG42709*) or haemocytes (*CG7882, pgant9, eya* and *CG30345*). Interestingly, *moody* is one of the most known markers of the *Drosophila* BBB glia and its ortholog in mammals (*Gpr84*) is a well-known pro-inflammatory maker that is highly up regulated in microglia upon nerve injury (76). Revisiting the role of *moody* during neuroinflammation could shed light into the function of this gene in both species.

As a corollary of the above and based on multiple pieces of evidence, we propose that *repo* constitutes the *bona fide* fly gliogenic gene. Accordingly, Repo misexpression in the mesoderm suppresses haematopoiesis and its lack triggers the expression of haemocyte markers in the nervous system (77). We speculate that the Gcm pathway has an ancestral, conserved, role is in immunity and may have been coopted in the differentiation of fly glia. One of the future challenges will be to characterize the Gcm-positive cells associated with the aged brain.

### The conserved role of the Gcm pathway in immune processes

Together with the inhibitory role of Gcm on the JAK/STAT pathway and the increased response to wasp infestation, which relies on the Toll cascade, the present data strongly suggest that Gcm controls different inflammatory conditions, including aging (12). This is also in line with the transcriptomic data of the *gcmKD* animals, which confirm the induction of the JAK/STAT pathway observed *in vivo* and extend the inhibitory role of Gcm to other pathways such as IMD. Furthermore, *gcm* GOF can ameliorate the inflammatory phenotype in flies, thus paving the way for new therapeutic strategies against autoimmune diseases, such as MS, where the inflammatory response needs to be contained. We hypothesise that the induction of the Gcm cascade reduces the intensity of the inflammatory response and hence has a protective function. This hypothesis is corroborated by recent data obtained in other organisms. A peculiar macrophage population called pigment cells is present in the sea urchin (78). Such cells are involved in the immune defence by the production of a pigment that has anti-microbial properties. Morpholino antisense oligonucleotides for *Spgcm* (*gcmMO*) injection showed that *gcmMO* animals are less resistant to challenging environmental conditions portrayed by decreased survival rate compared to the control (79–81). A recent study showed that a *gcm* ortholog is also expressed in glia-like cells of the freshwater crayfish (*Pacifastacus leniusculus*) upon an acute inflammatory response (82). Moreover, the expression of a planaria *gcm* ortholog was found to be induced upon regeneration in a subset of cells close to the wound; its silencing has no effect in homeostatic conditions but impairs neoblast repopulation upon wounding (83, 84). Thus, more and more studies highlight an evolutionarily conserved mechanism.

In sum, we report here the discovery of an anti-inflammatory transcriptional cascade that is conserved from flies to humans. Given the strong potential of transcription factors in coordinating the expression of several genes and the scarce number of known transcription factors with a similar function, this work represents a major contribution to understand the molecular mechanisms controlling the inflammatory response. It also lays the ground for studying novel therapeutical targets for neuro-inflammatory diseases in humans.

## Author contributions

AP, SM, PC, RP, BNO and AG designed the experiments and co-wrote the manuscript. AP, SM, CR, RP performed the experiments in mice, RP performed the experiments in humans, SM and AP performed the assays in *Drosophila melanogaster*, YY was responsible for the mGcm2^flox/flox^ production and PC analysed the RNAseq data.

## Acknowledgements

We thank the Imaging Center of the IGBMC for technical assistance, as well as the Mouse Clinic (ICS) for producing of the mGcm2^flox/flox^ strain. We also thank the ICM mouse facility (ICMice), the ICM histology (Histomics) and the cellular imaging (ICM-Quant) facilities. AP was supported by the ARSEP Foundation and the grant from Laboratoires d’excellence (LabEx INRT), SM by CEFIPRA and the FRM foundations, YY by the ARSEP foundation. RP was funded by the ARSEP foundation and NeurATRIS.

This work was supported by INSERM, CNRS, UDS, Ligue Régionale contre le Cancer, Hôpital de Strasbourg, ARC, CEFIPRA, ANR grants, the CNRS/University LIA Calim, The French MS foundation ARSEP, the Investissements d’Avenir ANR-10-IAIHU-06 (IHU-A-ICM) and ANR-11-INBS-0011 (NeurATRIS). The IGBMC was also supported by a French state fund through the ANR labex. We are grateful to the UK MS tissue Bank (Imperial College, London, UK) for providing post-mortem MS brain samples and to C. Linnington and I. Ando for providing antibodies.

## Notes

### Competing Interest Statement

The authors have declared no competing interest.

