## Supplemental data for "Gcm: a novel anti-inflammatory transcriptional cascade conserved from flies to humans"

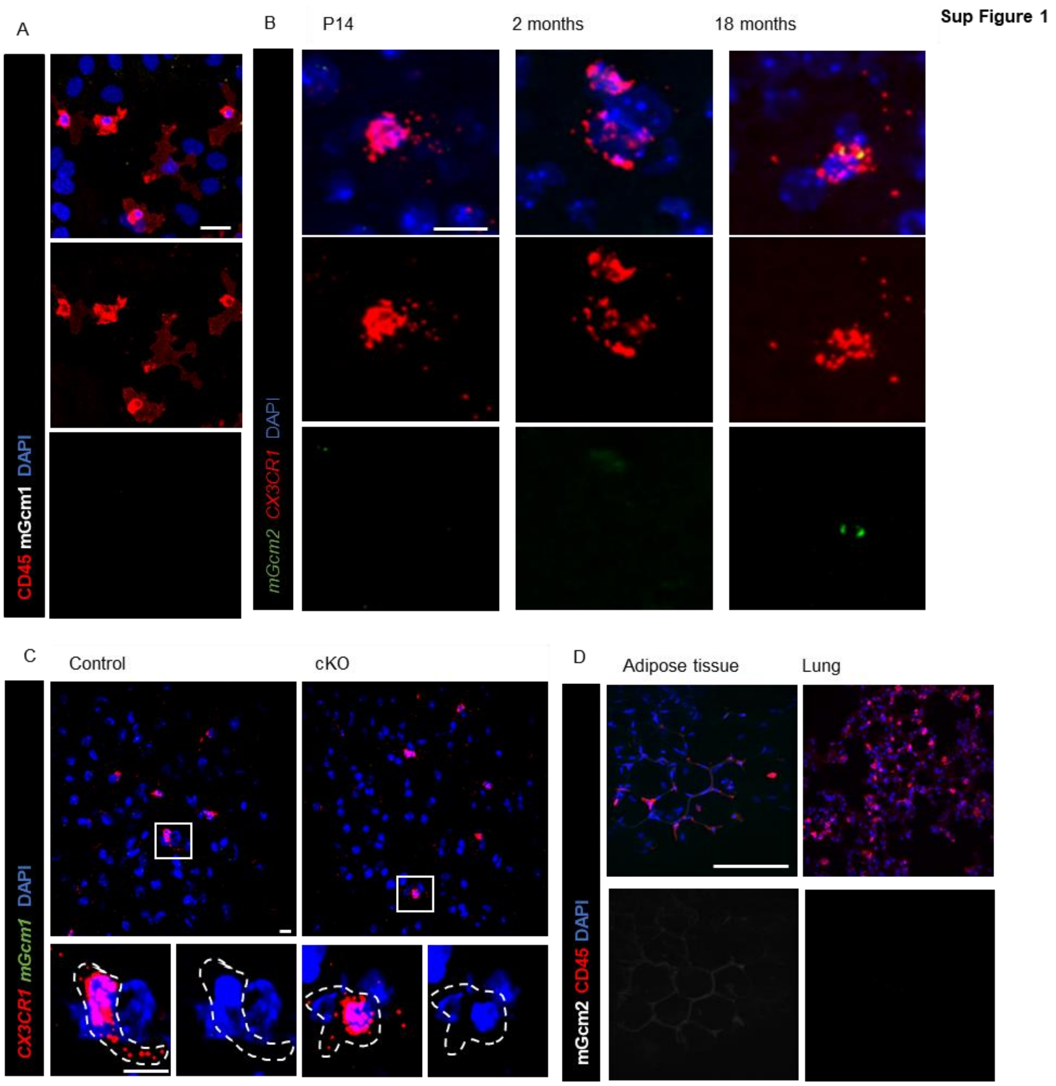


**Supplementary figure 1**. **mGcm1 and mGcm2 expression**. (**A**) Immunolabelling of CNS primary cultures for mGcm1 (grey), microglia (CD45, red) and DAPI (blue), N=3; scale bar 10 µm. (**B**) RNA ISH of brain sections on P14, 2 and 18month-old brain sections from control and cko animals: *Cx3cr1* (red), *mGcm2* (green) and DAPI (blue), N=3; scale bar: 10 µm. (**C**) RNAscope labelling on control and cKO brain sections at 18 months: *Cx3cr1* (red), *mGcm1* (green), DAPI (blue), N=3; scale bar: 10 µm. (**D**) Immunolabelling of resident macrophages from lung and adipose tissues at 24month-old animals: mGcm2 (grey), CD45 (red) and DAPI (blue). Scale bar: 100 µm. *Cx3cr1-Cre*$^{+/-}$;*mGcm2*$^{flox/}$ (control) and *Cx3cr1-Cre*$^{+/-}$, *mGcm2*$^{flox/flox}$(cKO)

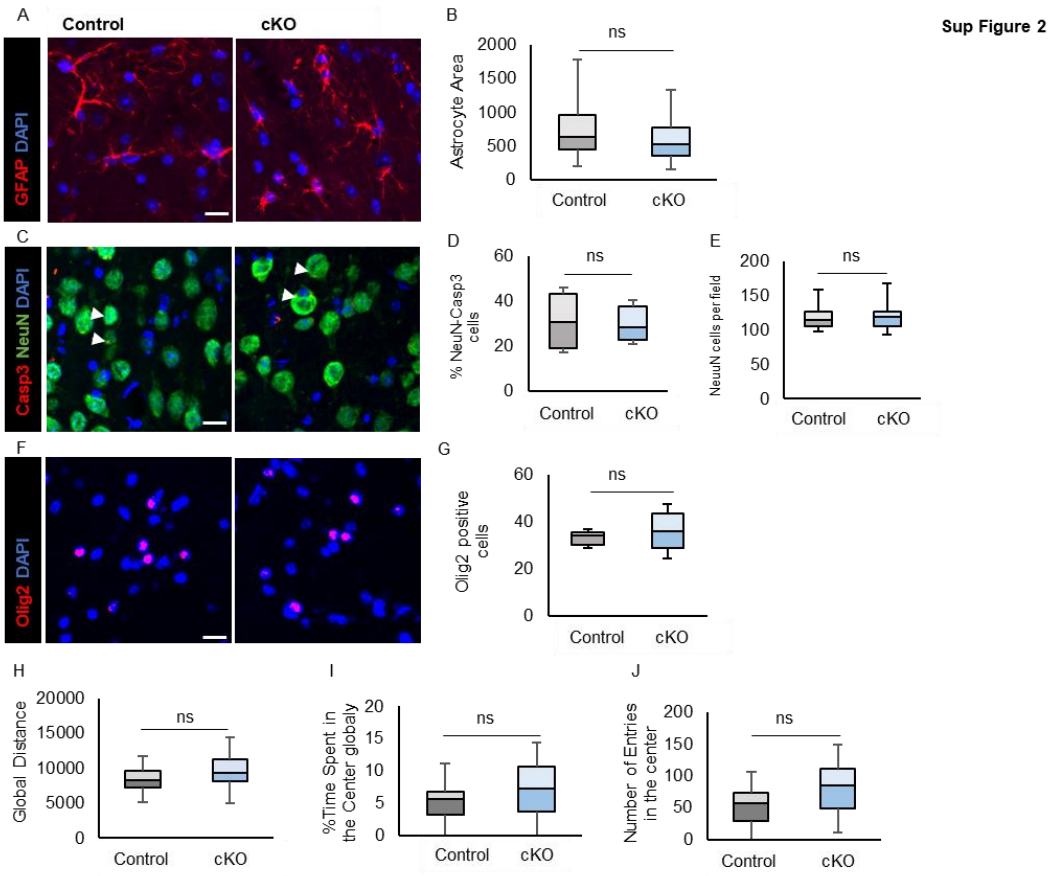


**Supplementary figure 2. Characterisation of cell populations in the cortex of 24month-old animals**. (**A**) Immunolabelling of astrocytes from control and cKO animals for GFAP (red). (**B**) Quantification of the GFAP area. (**C**) Immunolabelling of neurons with the pan neuronal marker NeuN (green) and the cell death marker Caspase3 (red). (**D**) Quantification for double positive NeuN-Caspase3 cells to calculate the neuronal cell death and (**E**) the total neuronal number. (**F**) Immunolabelling for oligodendrocytes with Olig2 and (G) quantification of oligodendrocytes in control and cKO animals. (**H,J**) Open field test for control and cKO animals at 18 months. The animals were evaluated for the total distance covered (**H**), the percentage of time spent in the centre (**I**) and how many times they entered in the centre (**J**). p-value: *<0.05, **<0.01, ***<0.001, and ns for not significant. N=5-13; scale bar: 50 µm. Statistical significance was determined by one-way ANOVA followed 2-tailed, unpaired t-test. *Cx3cr1-Cre*$^{+/-}$;*mGcm2*$^{flox/}$ (control) and *Cx3cr1-Cre*$^{+/-}$, *mGcm2*$^{flox/flox}$(cKO)

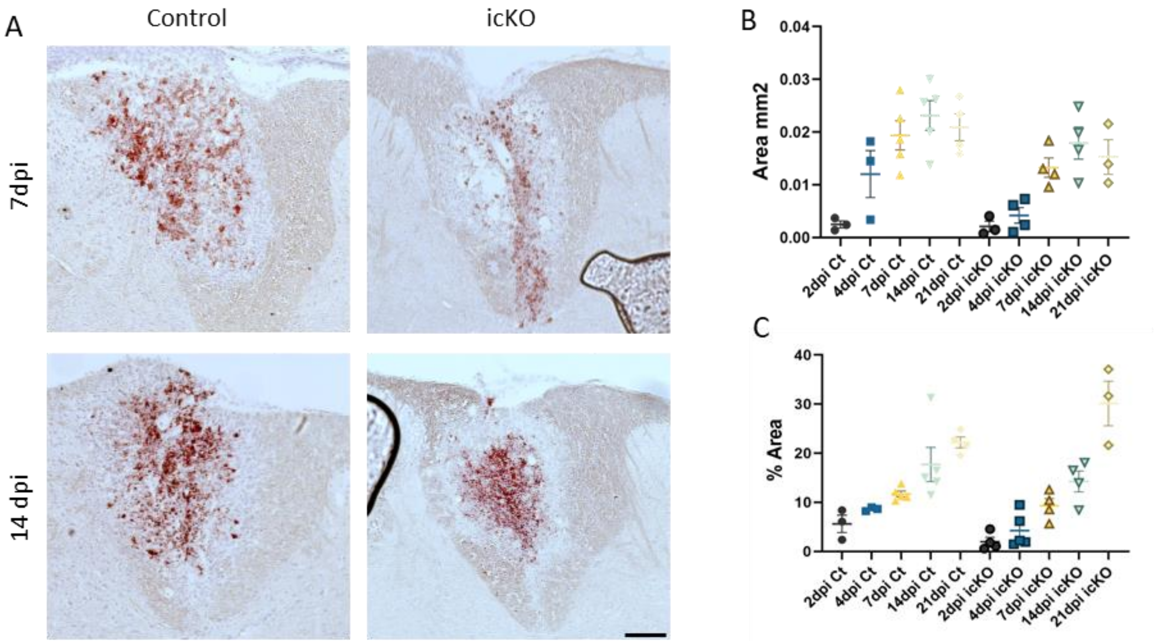


**Supplementary Figure 3. The density of macrophages containing myelin debris is not altered in LPC demyelinated lesions of *mGcm2* icKO mice. (A)** Oil-Red O staining of macrophages with myelin debris in control and *mGcm2* icKO LPC lesions, at 7 and 14 dpi. (**B**) Quantification of Oil-Red-O+ area in LPC lesions from 2 to 21 dpi, in control and *mGcm2* icKO animals. (**C**) Percentage of Oil-Red-O+ area in LPC lesions from 2 to 21 dpi in both experimental groups. Note that the clearance of myelin debris by macrophages in demyelinated lesions is not altered in *mGcm2* icKO with respect to control mice. Scale bar: (**A**), 100µm.

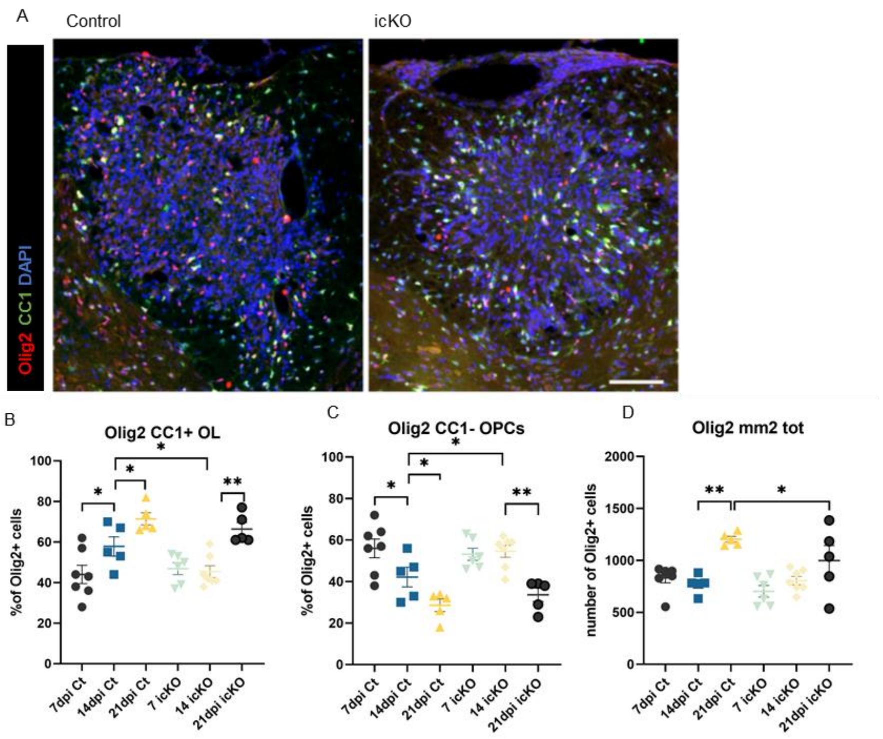


**Supplementary Figure 4. Loss of *mGcm2* function in microglia delays oligodendrocyte differentiation in demyelinated lesions.** (**A**) Immunolabelling with Olig2 (red) and CC1 (green) in LPC lesions of the spinal cord at 14 dpi, in control and *mGcm2* icKO mice. Nuclei are counterstained with DAPI. (**B,C**) Graphs indicating the percentage of Olig2/CC1-double positive differentiated oligodendrocytes and Olig2-positive/CC1-negative OPCs in demyelinated lesions from 7 to 21 dpi, in control and *mGcm2* icKO mice. (**D**) Quantification of the number of total Olig2-positive oligodendroglia in LPC lesions at 7, 14 and 21 dpi, in control and *mGcm2* icKO animals. Two-way ANOVA followed by Tukey's multiple comparison tests were used for statistical analysis. *p< 0.05, **p< 0.01. Scale bar: (**A**), 100 µm.

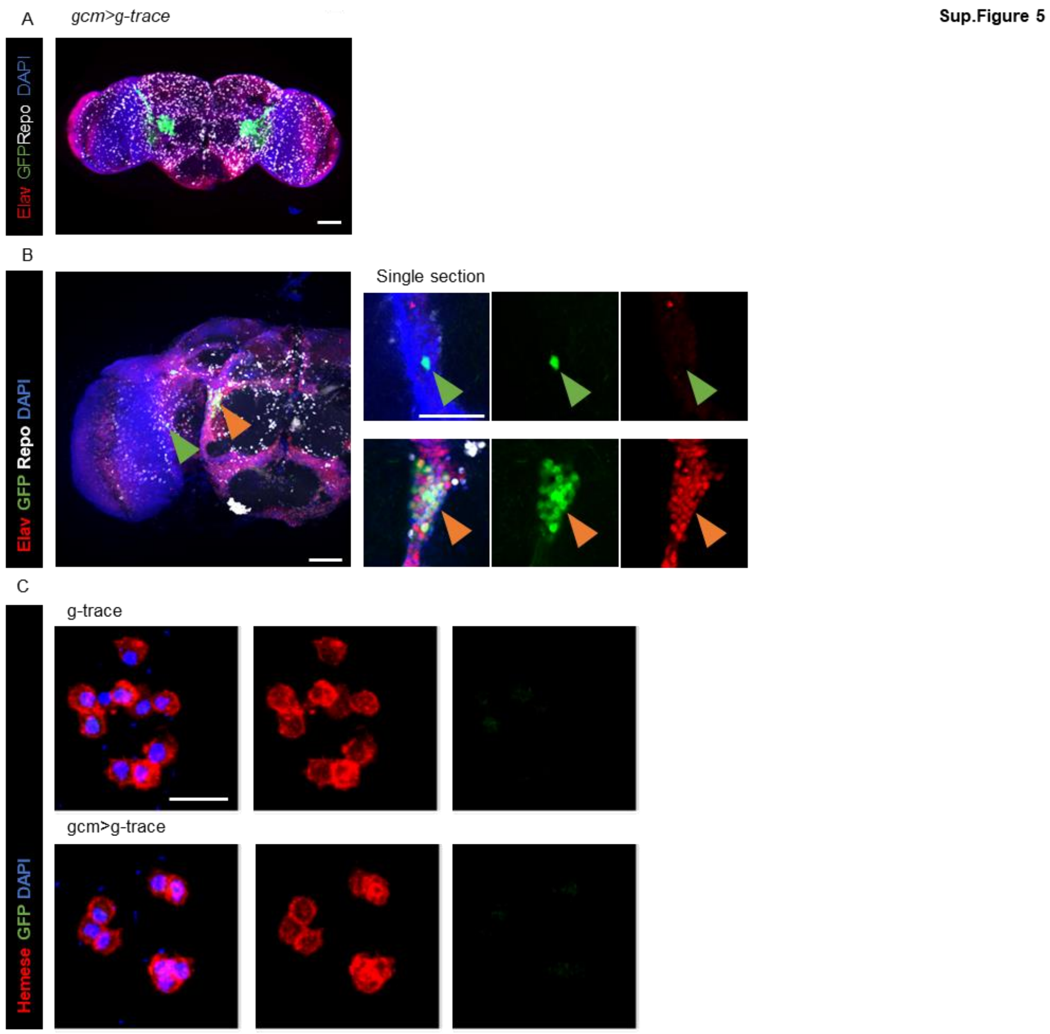


**Supplementary Figure 5. An inducible system tracing *gcm* expression**. (**A**) Immunolabelling of adult *Drosophila* brain with *gcm* tracing (*gcm>g-trace*=GFP) in green, Repo (white), Elav (white) and DAPI (blue). (**B**) Immunolabelling of adult *Drosophila* brain with *gcm* tracing (*gcm>g-trace*) in green, Repo (grey), Elav (red) and DAPI (blue), Scale bar: 20 µm. (C) Immunolabelling of hemocytes from gcm>g-trace strain. The pan-hemocyte marker Hemese (red), g-trace (green), DAPI (blue). Animals were raised at 18°C and shifted at 29°C at second instar larval. scale bar: 20 µm.

| Host | Target | Dilution | Reference |
|---|---|---|---|
| mouse | anti-Arginase1 | 1/200 | sc-271430 |
| rat | anti-CD11b | 1/400 | MCA-74G |
| rat | anti-CD45 | 1/50 | MCD-4500 |
| rat | anti-CD68 | 1/400 | MCA-1957 |
| rat | anti-F4/80 | 1/100 | MCA-497 |
| rabbit | anti-Gcm1 | 1/50 | B. Nait-Oumesmar |
| rabbit | anti-Gcm2 | 1/200 | ab201170 |
| rabbit | anti-Iba1 | 1/300 | 1919741 |
| mouse | anti-iNOS | 1/100 | ab210823 |
| mouse | anti-GFAP | 1/100 | PCN |
| mouse | anti-NeuN | 1/100 | ab279296 |
| rabbit | anti-Casp3 | 1/200 | mAb #9664 |
| mouse | anti-MOG | 1/20 | clone C18C5, C. Linnington |
| rabbit | anti-Olig2 | 1/400 | AB9610 |
| mouse | anti-APC | 1/100 | OP80 |
| mouse | anti-HLA | 1/100 | CR3/43 |
| mouse | anti-Repo | 1/20 | 8D12 (DSHB) |
| rat | anti-Elav | 1/100 | 7E8A10 (DSHB) |
| mouse | anti-Hemese | 1/40 | I. Ando |
| mouse | anti-L4 | 1/50 | I. Ando |
| chicken | anti-GFP | 1/500 | 703095155 |
| donkey | anti-mouse-Cy3 | 1/500 | 715-165-151 |
| donkey | anti-rabbit-Cy3 | 1/500 | 711-165-152 |
| goat | anti-rat-Cy3 | 1/500 | 112-165-167 |
| goat | anti-mouse-Alexa647 | 1/500 | 115-605-166 |
| goat | anti-rabbit-Cy5 | 1/500 | 111-175-144 |
| goat | anti-rat-Cy5 | 1/500 | 112-175-144 |
| donkey | anti-chicken-FITC | 1/500 | A11039 |
| goat | anti-rat-Alexa568 | 1/1000 | A11077 |
| donkey | anti-rat-Alexa488 | 1/1000 | A21208 |
| donkey | anti-rat-Alexa647 | 1/1000 | AB150155 |
| donkey | anti-rabbit-Alexa568 | 1/1000 | A10042 |
| goat | anti-mouse-Alexa488 | 1/1000 | 1070-02 |
| donkey | anti-mouse-Alexa488 | 1/1000 | A21202 |
| goat | anti-mouse-FITC | 1/100 | A21141 |

**Supplementary Table 1. Primary and secondary antibodies**

| **MS cases** | **Age** | **Sex** | **PMD** | **Disease duration** | **Disease course** | **Lesions analyzed** | | | |
|---|---|---|---|---|---|---|---|---|---|
| | | | | | | Active | Chronic active | Chronic inactive | Shadow plaques |
| MS94CLA6 | 42 | F | 11 | 6 | PP | **1** | | | |
| MS100CLC2 | 46 | M | 7 | 8 | SP | | | **1** | |
| MS121CLC5 | 49 | F | 24 | 14 | PR | **1** | | **1** | |
| MS106L6C8 | 39 | F | 18 | 21 | PP | | **1** | | |
| **Control** | | | | | | | | | |
| C26CLA5 | 79 | F | 18 | Cardiac failure | | | | | |

**Supplementary Table 2. MS and control cases used to study hGCM2 expression**

| | Target | Taqman probes |
|---|---|---|
| | Hprt | HPRT TaqMan® Gene Expression Assays (Mm03024075_m1) |
| | mGcm2 | GCM2 TaqMan® Gene Expression Assays (Mm00492312_m1) |
| M1 | iNOS | NOS2 TaqMan® Gene Expression Assays (Mm00440502_m1) |
| M1 | TLR2 | TLR2 TaqMan® Gene Expression Assays (Mm00442346_m1) |
| M2a | Arg-1 | ARG1 TaqMan® Gene Expression Assays (Mm00475988_m1) |
| M2b | Il-4ra | IL-4a TaqMan® Gene Expression Assays (Mm01275139_m1) |
| M2c | CD163 | Cd163 TaqMan® Gene Expression Assays (Mm00474091_m1) |

**Supplementary Table 3. Taqman probes used for qRT-PCR analysis of LPC lesions**
